# A temporal atlas of TDP-43 interactions reveals persistent network remodelling beyond stress granule clearance

**DOI:** 10.64898/2026.11.16.766431

**Authors:** Jerneja Nimac, Shujing Li, Luca Biasetti, Britta Eggers, Karin Schork, Helena Motaln, Ana Marija Kodra, Klementina Polanec, Urša Čerček, Klara Razboršek Fink, Florencia Vassallu, Lucas Muzio, Bradley N Smith, Vera Župunski, Martin Eisenacher, Katrin Marcus, Lionel Muller Igaz, Jernej Ule, Claire Troakes, Boris Rogelj

**Affiliations:** Jozef Stefan Institute, Department of Biotechnology, 1000 Ljubljana, Slovenia; Graduate School of Biomedicine, Faculty of Medicine, University of Ljubljana, Ljubljana, 1000, Slovenia; Department of Basic and Clinical Neuroscience, Maurice Wohl Clinical Neuroscience Institute, Institute of Psychiatry, Psychology & Neuroscience, King’s College London, London SE5 9RX, United Kingdom; London Neurodegenerative Diseases Brain Bank, Department of Basic and Clinical Neuroscience, Institute of Psychiatry, Psychology & Neuroscience, King’s College London, London, United Kingdom; Medizinisches Proteom-Center, Medical Faculty, Ruhr-University Bochum, Bochum, Germany; Medical Proteome Analysis, Center for Protein Diagnostics (PRODI), Ruhr-University Bochum, Bochum, Germany; Core Unit for Bioinformatics (CUBiMed.RUB), Medical Faculty, Ruhr-University Bochum, Bochum, Germany; BioInfra.Prot, Deutsches Netzwerk für Bioinformatik-Infrastruktur - de.NBI, Medical Faculty, Ruhr University Bochum, Bochum, Germany; University of Ljubljana, Faculty of Chemistry and Chemical Technology, Ljubljana, Slovenia; Universidad de Buenos Aires, Facultad de Ciencias Médicas, Departamento de Ciencias Fisiológicas. Grupo de Neurociencias de Sistemas. Buenos Aires, Argentina; CONICET - Universidad de Buenos Aires. Instituto de Fisiología y Biofísica Bernardo Houssay (IFIBIO Houssay). Buenos Aires, Argentina; UK Dementia Research Institute at King’s College London, 5 Cutcombe Rd, London, UK

**Keywords:** TDP-43, dynamic protein interaction network, proximity proteomics, APEX2, stress granules, stress response, stress recovery, ALS, FTD

## Abstract

Cytoplasmic aggregation of the nuclear protein TDP-43 is associated with several neurodegenerative disorders and may be promoted by cellular stress. However, the dynamics of TDP-43 protein−protein interactions during stress and recovery remain poorly understood. Here, we combined APEX2 proximity labelling with quantitative mass spectrometry to generate a temporal atlas of wild-type TDP-43 and pathology-mimicking TDP-43dNLS and TDP-43CTF variants. Interactomes were profiled across steady state, oxidative stress, and recovery. We reveal extensive state- and variant-dependent remodelling of TDP-43 interaction networks, uncovering distinct temporal interaction patterns throughout the stress response and recovery. Notably, distinct TDP-43 interaction signatures persisted after stress granule clearance, suggesting a molecular memory of stress. Together, our study establishes a comprehensive temporal resource for investigating TDP-43 interaction dynamics and provides a framework for understanding how stress-induced network alterations may contribute to TDP-43 proteinopathies.

## Introduction

Neurodegenerative disorders, such as Alzheimer’s disease (AD), Parkinson’s disease (PD), amyotrophic lateral sclerosis (ALS), and frontotemporal dementia (FTD), affect over 57 million people globally, with incidence continuing to rise^1^. These clinically heterogeneous diseases, characterised by progressive loss and abnormal function of neurons in the central or peripheral nervous system^2,3^, currently lack effective treatments^4^. Ageing, environmental stress, oxidative stress, and protein dysfunction are common risk factors for neurological disorders^5^, and one of the hallmarks of these disorders is the formation of insoluble protein aggregates in the central nervous system^6^. Transactive response DNA-binding protein of 43 kDa (TDP-43) forms toxic aggregates in ALS and FTD^7^, which are also frequently observed in other diseases, including AD, limbic-predominant age-related TDP-43 encephalopathy (LATE), and PD^6,8^. As TDP-43 is a central protein in the pathogenesis of these diseases, they are often referred to as TDP-43 proteinopathies^9,10^.

TDP-43 shuttles between the nucleus and cytoplasm, where it plays roles in various cellular processes, including RNA metabolism, transcription regulation^11^, nuclear transport^12,13^, alternative splicing^14^, mRNA expression, and mRNA stability^11,15^, each of which is fine-tuned by its interactions with other molecules, particularly RNA and proteins^9^. Disrupted or altered dynamics of protein interactions with TDP-43 may therefore contribute to its accumulation in disease^9^. These protein interactions can be irreversibly altered under different conditions, such as cellular stress, especially as stress response mechanisms are impaired in neurodegeneration^9,16,17^. One such mechanism is the formation of stress granules (SGs), cytoplasmic membrane-less organelles to which TDP-43 can be recruited under stress^16,18,19^. Under normal conditions, SGs facilitate cell survival by storing mRNAs, translation factors, and RNA-binding proteins, and are disassembled as the cell recovers^18^. However, prolonged cellular stress, together with ineffective stress responses, can create an intracellular environment that results in aggregation of SG components, including TDP-43^18,19^. While the TDP-43 interactome under physiological conditions and stress has been extensively studied^20–22^, the interactome during recovery, specifically during SG disassembly, remains unknown.

In this study, we used the APEX2 proximity labelling method^23,24^ combined with mass spectrometry to identify dynamic TDP-43 interactomes under steady state, oxidative stress, mid-recovery, and full post-stress recovery conditions. Using this approach, we mapped the proximal interactome of wild-type TDP-43 (TDP-43wt), the 25 kDa C-terminal fragment of TDP-43 (TDP-43CTF) found in pathological aggregates, and TDP-43 lacking a nuclear localization signal (TDP-43dNLS), which mimics pathological conditions. We classified interactors into clusters based on their interaction profiles with each TDP-43 variant from steady state to full recovery. We identified interaction trajectories of previously established interactors of TDP-43, including hnRNPA2B1, KPNA3 and VIM, as well as novel interactors, where we focused on YTHDF3. This systematic study revealed altered interaction behaviour between some interactors and TDP-43 variants across steady state, stress and recovery states, and identified new proteins involved in neurodegeneration, which we also investigated in pathology. Our study provides valuable resources on the dynamic interactomes of TDP-43 and its pathology-mimicking variants during stress and recovery in physiological and disease conditions, and offers novel insights into the molecular memory mechanisms of stress response proteins that could guide future aggregation studies.

## Results

### Determination of the proximal interactome of TDP-43 under stress and recovery

To investigate the interactome of TDP-43 under oxidative stress, during post-stress recovery, and potentially in pathology, we used the APEX2 method^23^ (Figures 1A). The interactomes of TDP-43wt, TDP-43dNLS, which mimics disease conditions, and TDP-43CTF, found in pathological aggregates, were determined. To minimize non-specific labelling and toxic effects of overexpressed proteins, inducible stable HEK293 Flp-In cell lines were generated (Figures 1A−1C). Each bait protein, together with its corresponding interactors, exhibited the expected subcellular localization, whereas bait APEX2-myc, used as a control, was present in all cellular compartments (Figures 1B, S1A and S1B). Validated stable cell lines were then subjected to the APEX2 method, and samples collected in the pull-down assay were analysed by mass spectrometry (MS) (Methods, Figures 1A and S1C–S1F). We determined the interactome for all TDP-43 variants and the control at four different time points: before oxidative stress (unstressed, 0 h), under oxidative stress (100% SG, 1.5 h), when the burden of stress granules in the cell was 50% (50% SG, 4 h) after oxidative stress removal, and 24 hours post-stress recovery, when the cell has returned to a stable state SG burden (recovery, 24 h) (Figures 1D, 2A and S2A–S2C).

**Figure 1:**
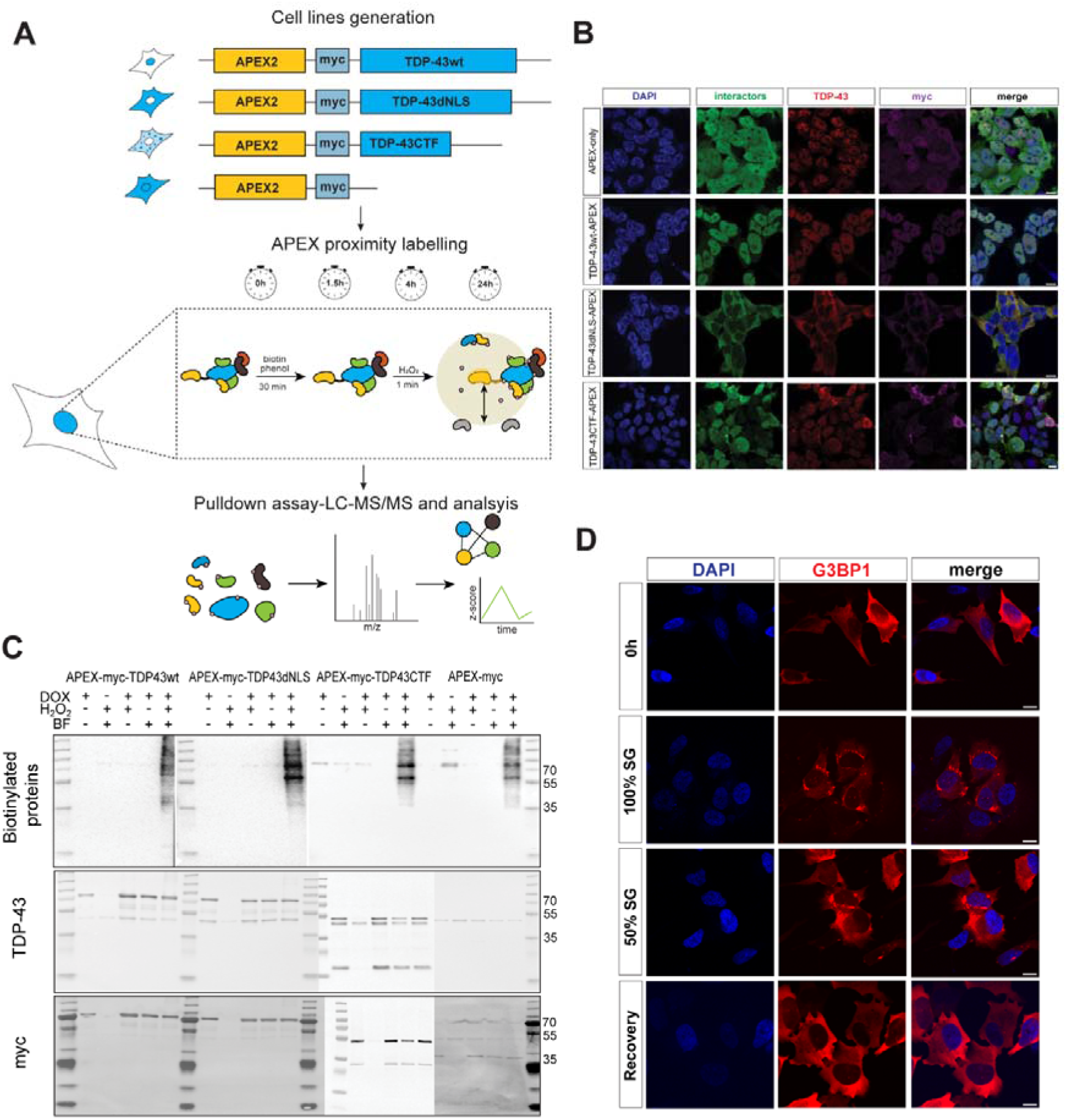
Profiling proximal proteomes of TDP-43wt, TDP-43dNLS, and TDP-43CTF under unstressed conditions, during oxidative stress, and during recovery. **(A)** Schematic overview of the experimental steps. Three variants of TDP-43, namely TDP-43wt, TDP-43dNLS and TDP-43CTF, fused with peroxidase APEX2, were stably expressed in HEK293 cell lines. APEX2-myc served as a control. Cells were then subjected to biotin phenol for 30 min and H_2_O_2_ for 1 min for labelling in triplicate. Cells were harvested at 0 h (unstressed), 1.5 h (during oxidative stress), 4 h (50% stress granule clearance), and 24 h (full post-stress recovery) and biotinylated interactors were isolated and identified with mass spectrometry analysis. (B) Immunofluorescence images show that TDP-43wt was mainly present in the nucleus, TDP-43dNLS in the cytoplasm, and TDP-43CTF formed aggregates.

**Figure 2:**
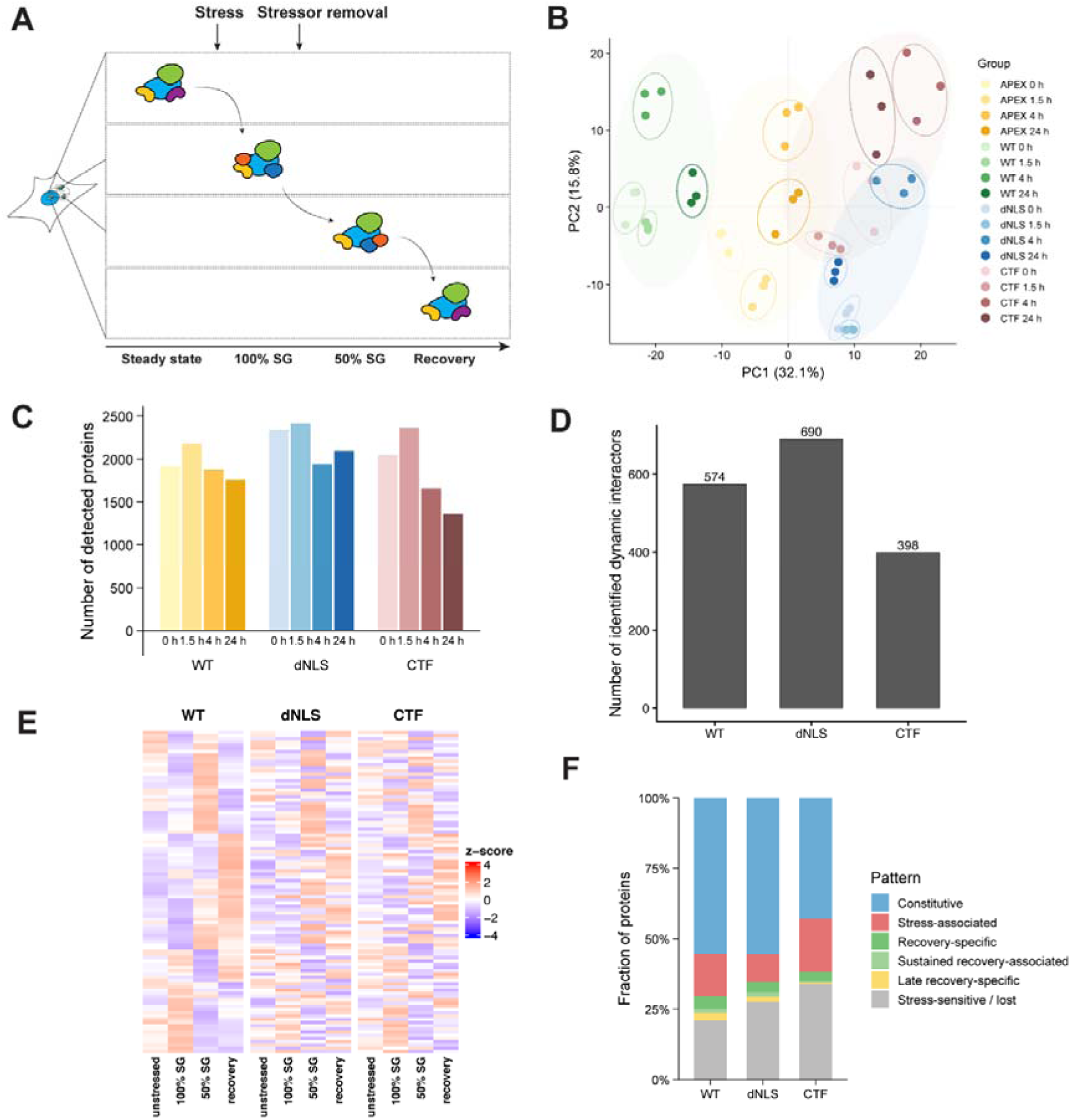
Global overview and classification of the dynamic interactome. **(A)** Schematic overview of dynamic changes in the TDP-43 interactome over time (unstressed, during oxidative stress, and during mid and late recovery). Arrows indicate stress stimuli and stressor withdrawal. **(B)** Principal component analysis (PCA) graph showing separation of replicates and grouping of samples based on the time point at which the sample was harvested and the construct present. The plot shows similar global profiles for pathology-mimic TDP-43 variants. Bar plots present **(C)** the number of detected proteins per sample group, where a protein was detected in at least 2 out of 3 replicates, and **(D)** the number of proteins classified as dynamic interactors across the stress-recovery time course. These proteins had an ANOVA FDR < 0.05, no fold-change cutoff was applied. **(E)** Heatmap of 500 proteins associated with TDP-43wt, TDP-43dNLS, and TDP-43CTF dynamic interactome under four conditions (unstressed, 100% SG, 50% SG, and recovery) based on z-score-normalized intensities. Rows were ordered by hierarchical clustering of the WT dataset. The same order was applied to all constructs. **(F)** Using On/Off analysis, we classified proteins into biologically interpretable temporal categories based on their detection across the time course. Detection across the time course was converted into a binary On/Off pattern, a protein was considered detected in a condition if it was identified in at least 2 out of 3 replicates (more in Table S1).

The localization of interactors corresponds to the localization of the TDP-43 variant localization. HEK293 cells stably expressing APEX2-myc constructs were co-stained with anti-myc antibody (violet), anti-TDP-43 antibody (red), and streptavidin-Alexa Fluor 488 (green), and imaged by confocal microscopy. DNA was labelled with 4,6-diamidino-2-phenylindole (DAPI) (blue). **(C)** Western blot analysis of lysates after APEX method shows that fusion proteins were only expressed in the presence of DOX and were labelled in the presence of BF and H_2_O_2_. **(D)** Immunofluorescence images show the stress granule marker G3BP1 fused to the fluorescent protein mScarletI in SH-SY5Y Flp-In cells before oxidative stress (unstressed, 0 h), under oxidative stress (100% SG), after stressor removal when the SG burden had decreased by approximately 50% (50% SG), and after 24 h of recovery, when stress granules had largely resolved (recovery). mScarletI-G3BP1 is shown in red, images were acquired by confocal microscopy. DNA was labelled with 4,6-diamidino-2-phenylindole (DAPI) (blue). Scale bars, 10 µm. DOX – doxycycline, BF-biotin phenol.

After locally estimated scatterplot smoothing (LOESS) normalization, replicates of TDP-43 variants at different time points clustered more closely together. Samples of TDP-43 variants and control APEX2 were clearly separated by construct and by the time point at which the samples were harvested. Furthermore, TDP-43dNLS and TDP-43CTF variants were clustered closer together, showing similar global profiles (Figures 2B and S3A–S3D). The distribution of log2-transformed protein intensities was comparable across samples, suggesting consistent signal distribution and no major technical outliers (Figure S3E). In addition, the number of valid quantified proteins was also comparable across technical replicates (Figure S3F). Approximately 2,000 proteins were detected at each time point for TDP-43wt, TDP-43dNLS, and TDP-43 CTF after filtering for proteins quantified in at least two of three replicates (Figure 2C, Table S1). Among these proteins, 574 associated with APEX2-myc-TDP-43wt, 690 with APEX2-myc-TDP-43dNLS, and 398 with APEX2-myc-TDP-43CTF were classified as dynamic interactors, meaning these proteins had an ANOVA FDR < 0.05 across the stress-recovery time course (Figure 2D). To further profile interactor dynamics over time, clustering and On/Off analyses were performed (Figures 2E and 2F and Table S1). The heatmap revealed construct-specific changes in the TDP-43 interactome during stress and recovery (Figure 2E). Additionally, both analyses confirmed that a subset of interactors remains constitutively detected across all conditions, while some proteins display stress-associated, recovery-specific, or late-recovery-specific patterns. Together, these findings suggest dynamic remodelling of the TDP-43 interaction network over time (Figures 2E and 2F).

A Gene Ontology (GO) enrichment analysis of the significant interaction proteins (see Methods) for each TDP-43 variant and time point was also in good agreement with their expected localization, molecular function and biological process (Figures S4A–S4I, Table S2). Under stress, protein interactors associated with the stress response were mobilized to come into proximity with TDP-43, while during recovery protein interactors involved in translation, axon guidance, senescence, and chaperone activity were enriched in the TDP-43 interaction network (Figures S4A−S4I). Among the interactors of TDP-43 identified in our experiment, we found previously known interactors under both physiological and disease conditions^20–22,24,25^ (Table S1). Thus, we identified the stress granule protein G3BP1^24,26^ enriched in TDP-43wt during stress (Figure 5A), the chaperone protein HSPA9^27^ enriched in mid-recovery with all TDP-43 variants (Figure S5B), and the nuclear pore protein NUP153^20,28^ enriched with the ALS-associated variant TDP-43dNLS in the absence of oxidative stress, during stress, and in late recovery (Figure S5C). These findings indicate that the data quality meets expected standards. Moreover, our comprehensive analysis of the interactome of different TDP-43 variants during stress and recovery increases knowledge about TDP-43 interactors and serves as an in-depth resource for future studies of TDP-43 aggregation and proteins involved in the formation of pathological aggregates.

### Transient stress-related rewiring of the TDP-43 dynamic interactome

TDP-43 shuttles between the nucleus and cytoplasm under physiological conditions^29^. During cellular stress, the transfer of TDP-43 into the cytoplasm increases, where it is incorporated into stress granules with other proteins^30^. When the stress subsides, the stress granules dissolve, releasing TDP-43 and allowing its return to the nucleus^31^. During this process, the dynamics of TDP-43 protein interactions change. To evaluate these interaction changes, we captured snapshots of the TDP-43 interactome before stress, during stress, and during recovery after the stressor was removed. This enabled us to classify proteins into clusters based on their interaction trajectories over time. Proteins with similar interaction paths with individual TDP-43 variants were grouped into one cluster (Table S1). While some proteins followed the same interaction trajectory for all TDP-43 variants, for others this interaction differed between variants. In clusters containing SON, a nuclear speckle scaffolding protein^32^, interaction with TDP-43wt was enriched during stress and recovery (Figures 3A and 3G), whereas interaction with TDP-43dNLS increased over time (Figures 3B and 3H), and interaction with TDP-43CTF increased in mid-recovery (Figures 3C and 3I). A similar pattern was observed for the clusters containing hnRNPA2B1, a nuclear protein that binds m6A-modified RNA^33^ and is a known interactor of TDP-43. In the presence of TDP-43wt, interaction was enriched during stress and then decreased (Figures 3J and 3N), while in the presence of TDP-43dNLS, interaction increased over time, suggesting that this interaction is related to pathology (Figures 3K and 3O). Moreover, when comparing sets of proteins that interacted similarly within the same TDP-43 variant, we observed that these clusters differed among variants (Figures 3D–3F, 3L and 3M), supporting the hypothesis that a different interaction environment could also contribute to an ineffective response to stress and aggregation of TDP-43. Thus, interaction trajectories provide new insights into the dynamics of protein interactions with TDP-43 under physiological and pathological conditions, including the effects of TDP-43 mislocalisation and its presence in toxic aggregates.

**Figure 3:**
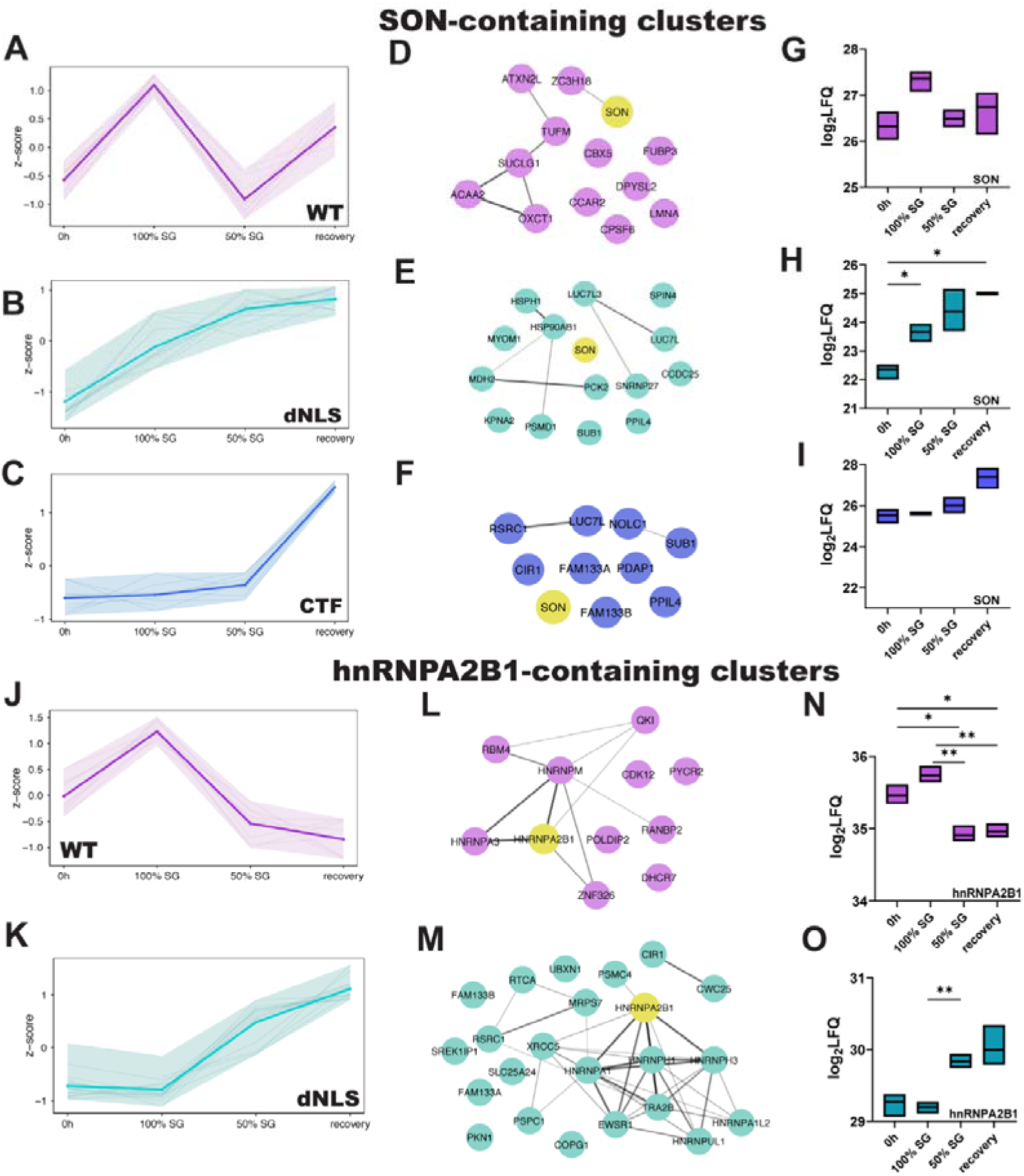
Dynamic interaction patterns of selected TDP-43 interactors during the stress response and protein cluster interaction networks that follow the same trajectory. (A-C) Interaction trajectory of SON-containing clusters with all three TDP-43 variants. Interaction with **(A)** TDP-43wt was enriched during stress, whereas interaction with **(B)** TDP-43dNLS increased across the time course, and for **(C)** TDP-43CTF, interaction increased during mid- and late-recovery. Each semi-transparent line represents the interaction behaviour of individual proteins of the cluster. The bold line represents the average trajectory, the shaded ribbon shows the full observed range, defined by minimum and maximum z-score values across all proteins at each time point. **(D-F)** Proximal protein cluster networks that followed the same trajectory differed among TDP-43 variants – **(D)** TDP-43wt cluster network, **(E)** TDP-43dNLS cluster network, and **(F)** TDP-43CTF cluster network. Nodes representing known interactors annotated in STRING are connected by edges. **(G-I)** Floating bar plots show log2-transformed LFQ intensity values of SON over the time course of interaction with **(G)** TDP-43wt, **(H)** TDP-43dNLS and **(I)** TDP-43CTF. Floating bars indicated the min to max range, with a line indicating the mean. Statistical analyses were performed using Welch-ANOVA, followed by a post-hoc test to compare time points in a pairwise manner, *p < 0.05, **p < 0.01, ***p < 0.001. **(J-K)** Interaction trajectory of hnRNPA2B1-containing clusters with TDP-43wt and TDP-43dNLS. TDP-43CTF did not interact with hnRNPA2B1. For TDP-43wt, interaction decreased after stressor removal, whereas interaction with TDP-43dNLS increased over time. Each semi-transparent line represents interaction behaviour of individual protein of the cluster. The bold line represents the average trajectory, the shaded ribbon shows the full observed range, defined by minimum and maximum z-score values across all proteins at each time point. **(L-M)** Proximal protein cluster network that followed the same trajectory differed among TDP-43 variants – **(L)** TDP-43wt cluster network, and **(M)** TDP-43dNLS cluster network. Nodes that represent known interactors annotated in STRING are connected by edges. **(N-O)** Floating bar plots show log2-transformed LFQ intensity values of hnRNPA2B1 over the time course of interaction with **(N)** TDP-43wt and **(O)** TDP-43dNLS. Floating bars indicated the min to max range, with a line indicating the mean. Statistical analyses were performed using Welch-ANOVA, followed by a post-hoc test to compare time points in a pairwise manner, *p < 0.05, **p < 0.01, ***p < 0.001.

### Molecular traces in the TDP-43 interactome linked to previous stress events

When comparing protein interaction trajectories among TDP-43 variants for different proteins, we discovered a stress memory mechanism that led to increased interactions during recovery, suggesting persistent interaction at later stages. When a cell is exposed to stress and the stressor is subsequently removed, the interactome is expected to return to its pre-stress state. However, after 24 h of recovery, when full recovery is anticipated, the interactome did not return to its initial state (Figure 4A). Traces of the previous stress persisted in the interactome, resembling a memory effect. Thus, in late recovery, TDP-43 gained new interactors and lost others (Figure 4B–4G). For example, KPNA3 interaction with TDP-43wt increased in late recovery (Figure 4D), as opposed the ceased interaction between VIM and TDP-43dNLS upon stress withdrawal (Figure 4G). To visually describe and interpret the enrichment analysis for these interactors of each TDP-43 variant, we grouped GO terms into functional and cellular compartment categories (Methods, Figures 5 and S6, Table S3). We observed that proteins classified as gained late recovery-associated interactors of TDP-43wt are part of stress-adaptive nuclear-cytoplasmic transport, proteostasis, and RNA-processing networks, while those lost in late recovery are involved in nucleolar and ribosome biogenesis, and RNA maturation processes. For TDP-43dNLS, the set of proteins gained in late recovery suggested a nuclear RNA-regulatory module enriched in RNA-binding, pre-mRNA splicing, transcriptional elongation, and cytoskeleton signalling. Compared with TDP-43wt, the TDP-43dNLS module appears more closely aligned with RNA-processing and nuclear body pathways highly relevant to TDP-43 proteinopathy.

**Figure 4:**
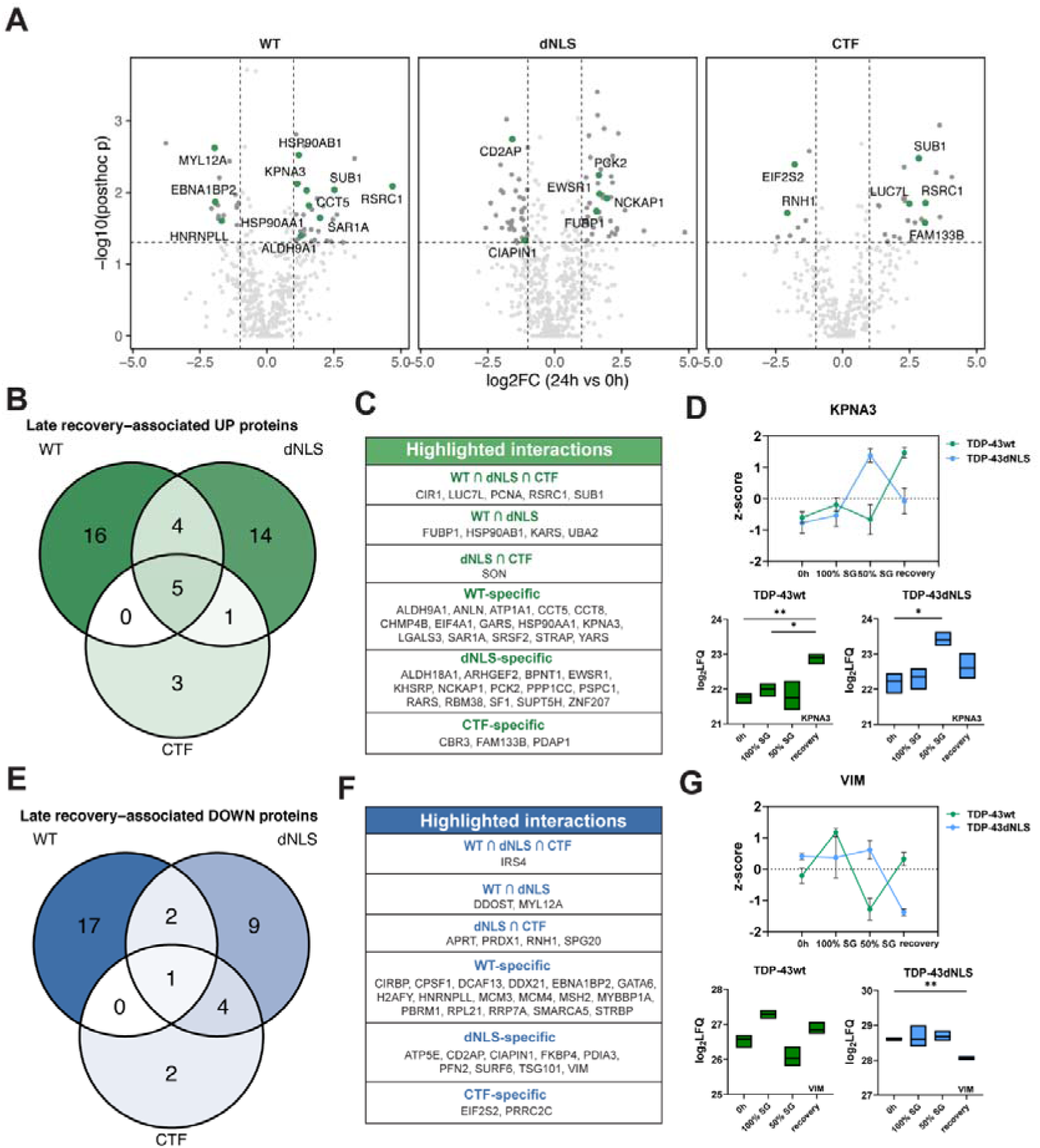
Late recovery-associated interactors of TDP-43 variants. **(A)** Volcano plots showing significantly upregulated and downregulated proximal interactors of TDP-43wt, TDP-43dNLS, and TDP-43CTF. The plots show relative protein levels at 24 h recovery compared to 0 h unstressed samples, with the x-axis on a log2 scale, analysed by MS. The y-axis shows the differential expression p-values on a -log10 scale. Green indicates late recovery-associated interactors with ANOVA FDR-adjusted p < 0.05, absolute FC ≥ 1.0. **(B)** Venn diagram of upregulated late recovery associated proteins. Proteins were included if they belonged to the late recovery-associated class and were detected in at least 2 of 3 replicates at all four time points. Proteins that passed the display-strict criteria in one construct were also included in another construct’s Venn set if that construct showed a recovery response in the same direction. **(C)** Table corresponding to the Venn diagram of upregulated late recovery-associated proteins. **(D)** KPNA3 interaction trajectory with TDP-43 variants over the time course, showing enrichment in late recovery with TDP-43wt. Data are presented as z-score mean ± SEM. The corresponding floating bar plot shows log2-transformed LFQ intensity values of KPNA3 over the time course of interaction with TDP-43wt and TDP-43dNLS. Floating bars indicated the min to max range, with a line indicating the mean. Statistical analyses were performed using Welch-ANOVA, followed by a post-hoc test to compare time points in a pairwise manner, *p < 0.05, **p < 0.01, ***p < 0.001. **(E)** Venn diagram of downregulated late recovery associated proteins. Proteins were included if they belonged to the late recovery-associated class and were detected in at least 2 of 3 replicates at all four time points, showed a negative recovery change. Proteins that passed the display-strict criteria in one construct were also included in another construct’s Venn set if that construct showed a similar recovery response in the same direction. **(F)** Table corresponding to the Venn diagram of downregulated late recovery-associated proteins. **(G)** VIM interaction trajectory with TDP-43 variants over the time course, showing loss of interaction in late recovery with TDP-43dNLS. Data are presented as z-score mean ± SEM. The corresponding floating bar plot shows log2-transformed LFQ intensity values of VIM over the time course of interaction with TDP-43wt and TDP-43dNLS. Floating bars indicated the min to max range, with a line indicating the mean. Statistical analyses were performed using Welch-ANOVA, followed by a post-hoc test to compare time points in a pairwise manner, *p < 0.05, **p < 0.01, ***p < 0.001.

**Figure 5:**
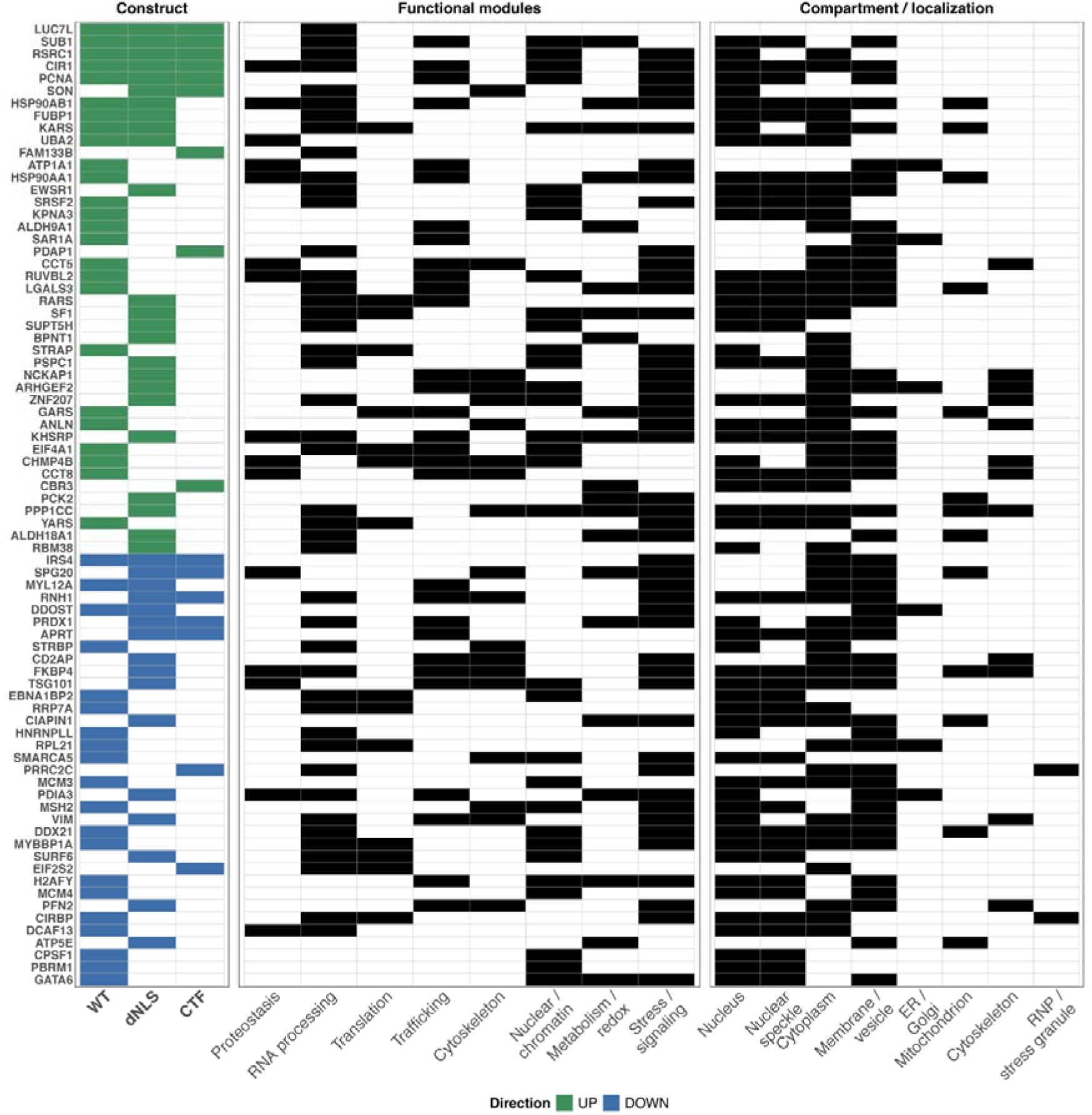
GO-based descriptive annotation profiling of late-recovery associated TDP-43 interactors. Upregulated and downregulated candidate proteins associated with late recovery for all constructs were annotated with functional and compartmental features. GO-based annotation panels were generated using descriptive keyword matching of GO terms retrieved through org. HS.eg. db and GO.db.

Moreover, under stress, TDP-43 protein variants recruited distinct sets of proteins, and some of these proteins continued to interact with TDP-43 variants as persistent post-stress interactors (Figures S5D– S5I). For instance, the interaction between SRSF3 and TDP-43CTF emerges after stress and increases over time (Figure S5F). In contrast, interactions between RPL6 and TDP-43 pathology mimicking variants decrease after oxidative stress, whereas a strong decrease is observed for TDP-43wt already during oxidative stress, followed by fluctuations during mid- and late-recovery (Figure S5I). Some stress-induced interactions that persist during recovery may help protect the cell from damage^34^, such as heat shock proteins that interact with misfolded proteins and remain in contact long after the stressor has been removed. This is important for maintaining protein homeostasis, supporting cell recovery after stress, and ensuring that misfolded proteins adopt the correct folding^35^. Thus, proteins that persist to interact with TDP-43wt after stress are involved in chaperone-mediated proteostasis, redox regulation, and homeostasis (Figures S5D, S5E, S6 and Table S3), whereas persistent post-stress proteins in the case of mislocalised TDP-43 forms are involved in RNA-processing, nuclear-cytoplasmic transport, and chromatin regulation (Figures S5D, S5E and 6, and Table S3). This suggests that TDP-43 mislocalisation can lead to altered nuclear transport and nuclear regulatory pathways, which could weaken the cellular response to stress.

These results provide a comprehensive characterization of the protein environments associated with different TDP-43 variants, the biological processes in which they participate, and the processes that are disrupted in the dNLS and CTF variants that mimic pathological conditions. We also identified proteins that remain in close proximity to TDP-43 or its variants following stress, as well as proteins that play important roles during the late stages of the recovery process.

### TDP-43 mislocalisation affects the m6A reader YTHDF3 in cellular model

To identify pathological TDP-43 interactors of potential clinical relevance, we examined the interactomes of TDP-43dNLS and TDP-43CTF across all time points. Analyses revealed several candidate interacting proteins, among which YTHDF3 was of particular interest. YTHDF3 is an m6A reader that recognizes m6A-modified RNAs, regulates their fate, and shuttles from the cytoplasm to the nucleus^36^. YTHDF3 also localizes to P-bodies, cytoplasmic stress granules, and neuronal RNA granules^37^. In addition to its other functions, YTHDF3 regulates the stability and translation of synaptic transcripts^38–40^. To validate its interaction with TDP-43 variants, we performed western blot analyses of proteins isolated by APEX2 proximity labelling (Figure 6A). These results, together with quantitative analysis of mass spectrometry data, showed that YTHDF3 preferentially interacted with TDP-43 variants that mimic disease conditions compared to the APEX2 control. The interaction was particularly enriched with TDP-43dNLS under all conditions, except during mid-recovery, when YTHDF3 interaction was similar for all TDP-43 variants compared to the control (Figure 6B). We further analysed z-scores and interaction trajectories, and observed that YTHDF3 is enriched during stress and becomes less abundant during recovery; however, this pattern was observed only with TDP-43dNLS (Figure 6C).

**Figure 6:**
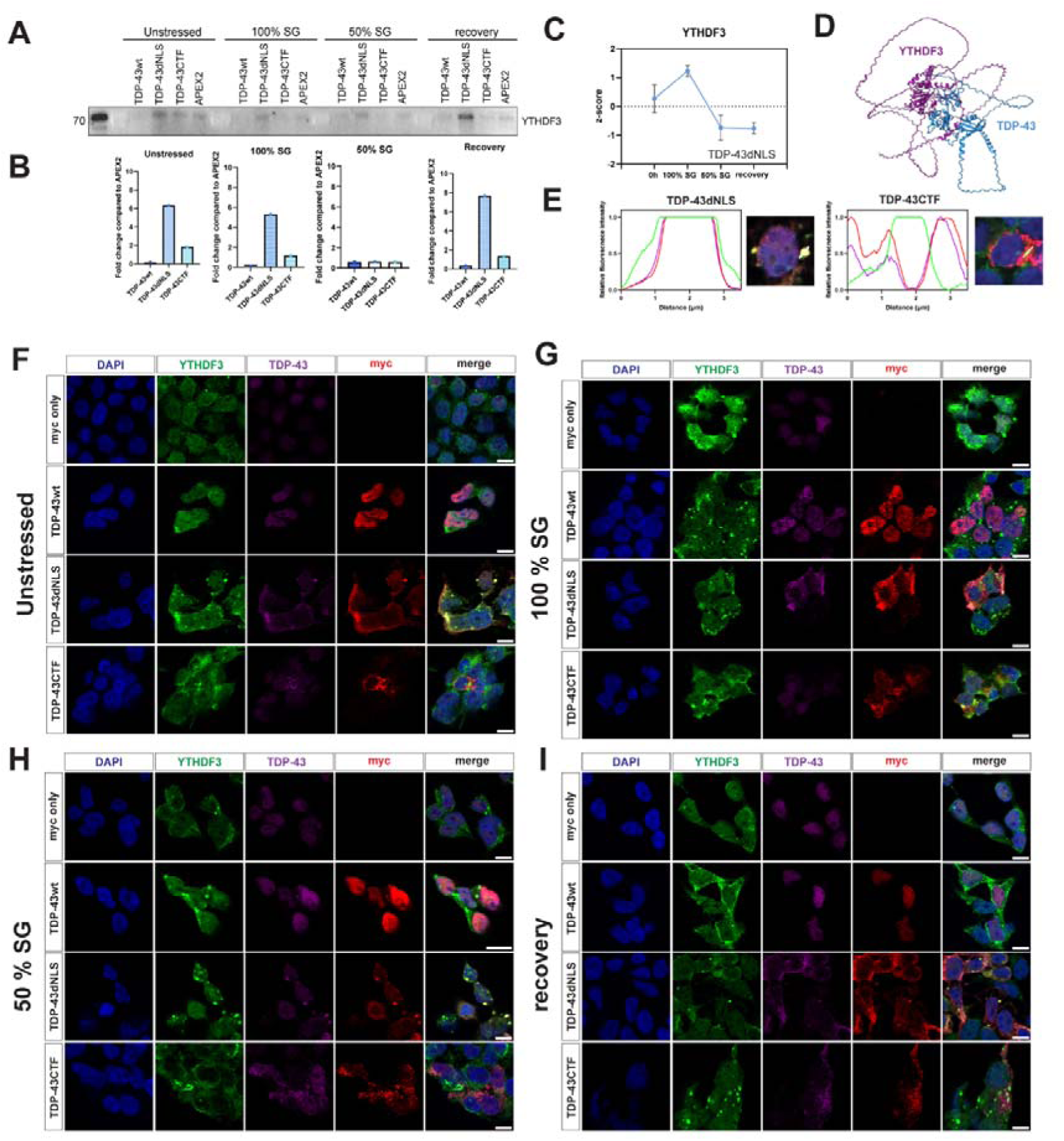
YTHDF3 granules colocalize with TDP-43dNLS granules. **(A)** Western blot analysis following the APEX2 method and streptavidin pull-down of biotinylated TDP-43 interactors for detection of YTHDF3. YTHDF3 was detected in unstressed, 100% SG, 50% SG, and recovery conditions. Experiment was repeated in three biological repeats. **(B)** Graphs showing MS quantification of YTHDF3. Results are presented as fold change of YTHDF3 in samples with TDP-43 variant compared to APEX2 control at each time point. **(C)** YTHDF3 interaction trajectory with TDP-43dNLS over the time course. Data are presented as z-score mean ± SEM. **(D)** AlphaFold model of TDP-43 and YTHDF3, showing interaction between YTHDF3 and the N-terminal region of TDP-43. **E-I,** Immunofluorescence images of TDP-43 variants and YTHDF3 behaviour over the time course. HEK293 cells expressing myc constructs were co-stained with anti-myc antibody (red), anti-TDP-43 antibody (violet), and anti-YTHDF3 antibody (green), and imaged by confocal microscopy. DNA was labelled with 4,6-diamidino-2-phenylindole (DAPI) (blue). Scale bars, 10 µm. **(E)** TDP-43dNLS granules colocalized with YTHDF3 granules, whereas TDP-43CTF aggregates surrounded YTHDF3 granules. Region of interest with accompanying line profile plots. Immunofluorescence in **(F)** unstressed cells, **(G)** stressed cells with 100% burden of stress granules, **(H)** mid-recovery when the burden of stress granules was 50%, and **(I)** expected full recovery after 24 h.

To visualise these changes in cells, we transfected HEK293 cells with myc-TDP-43wt, myc-TDP-43dNLS, myc-TDP-43CTF, and myc-only as a control. We then exposed these cells to oxidative stress for 90 min. Following this period, the culture medium was replaced for cells designated for recovery. Unstressed and stressed cells were then fixed, as were cells when the burden of stress granules was 50% and after full recovery (Figure 6F–I). We observed substantial overlap between YTHDF3 and TDP-43dNLS signals at all time points suggesting overlap between. In contrast overlap between YTHDF3 and TDP-43wt was only partial and mainly nuclear under steady state conditions (Figure 6F). In cells expressing TDP-43CTF, limited colocalization with YTHDF3 was observed during oxidative stress (Figure 6G). More intriguingly, we observed YTHDF3 to be surrounded by TDP-43CTF aggregates (Figure 6E), without directly colocalizing with them. This may be explained by the AlphaFold model, which suggests that YTHDF3 interacts with the N-terminal part of TDP-43 (Figure 6D). These findings also correlate with quantitative data from MS and western blot analysis (Figures 6A and 6B).

Since YTHDF3 is classified as a pre-stress seed, a protein with a high propensity to phase separate^24^, we next aimed to evaluate its effect on aggregation of a pathology-mimicking TDP-43 variant. For this reason, we co-transfected HEK293 cells with mKO2-TDP-43dNLS and EGFP-YTHDF3, stressed these cells, and followed TDP-43dNLS and YTHDF3 distribution (Figure S7). Using live imaging, we observed that TDP-43dNLS positive granules appeared simultaneously with YTHDF3-positive granules (Figure S7A), and both proteins seem to colocalize in these granules (Figure S7B), suggesting that TDP-43dNLS and YTHDF3 positive granules could represent seeds for pathological aggregation of TDP-43.

### TDP-43 pathology is linked with decreased YTHDF3 abundance

Proteins of the YTHDF family are important regulators of brain gene expression and may contribute to protein aggregation^38,39^. Our data from cellular models indicate that the interaction between a member of this family, YTHDF3, and TDP-43 could also be pathological. To investigate the clinical relevance of our finding in disease, we stained the brains of transgenic mice expressing TDP-43dNLS and control mice expressing TDP-43wt^41^. A clear decrease in YTHDF3 signal was observed in transgenic mice at the whole-neuron level and in both the nucleus and cytoplasm compared to controls (p < 0.0001, Figures 7A and 7B). However, we also observed that TDP-43dNLS mice have smaller neurons, including both nuclear and cytoplasmic compartments (Figures 7A and 7C). For this reason, we normalized the YTHDF3 signal by area, which showed that the decrease was preserved, suggesting that this is not the cause of the signal reduction (Figure S8A). Furthermore, when we silenced TDP-43 in HEK293 cells, we did not observe a change in YTHDF3 levels (Figure S8B), which is consistent with the data in neurons^42,43^. This evidence suggests that TDP-43 mislocalisation affects YTHDF3 reduction. Moreover, additional measurement of YTHDF3 subcellular distribution, in which we quantified the nuclear-to-cytoplasmic ratio, indicated a decrease in the nuclear/cytoplasmic YTHDF3 ratio in transgenic mice expressing TDP-43dNLS (Figure S8C and S8D), supporting a shift in YTHDF3 distribution from the nucleus to the cytoplasm.

**Figure 7:**
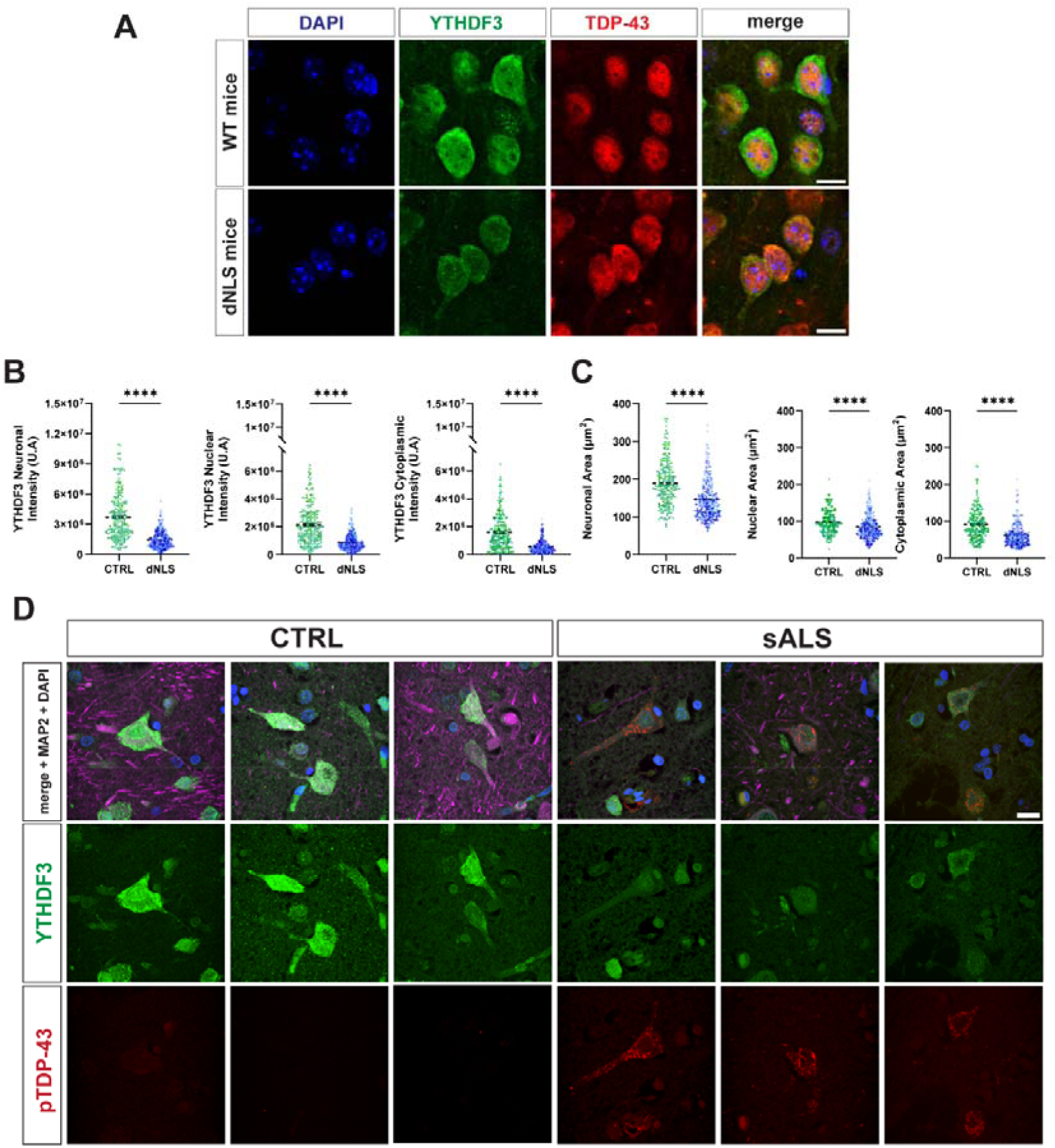
Interaction between TDP-43 and YTHDF3 contributes to ALS/FTD pathology. **(A)** Confocal images of mouse neurons from mice expressing TDP-43wt and transgenic mice expressing TDP-43dNLS, immunolabelled with anti-TDP-43 antibody (red) and anti-YTHDF3 antibody (green). DNA was labelled with 4,6-diamidino-2-phenylindole (DAPI) (blue). Scale bars, 10 µm. **(B)** Graphs showing quantitative analysis of YTHDF3 intensity in whole-neurons, nuclei, and cytoplasm in WT mouse neurons (n = 220) and transgenic TDP-43dNLS mouse neurons (n = 298). **(C)** Quantification of neuronal, nuclear, and cytoplasmic area in WT mouse neurons (n = 220) and transgenic TDP-43dNLS mouse neurons (n = 298). Brain tissue from four mice per group was analysed; different shades of blue and green correspond to different animals. Statistical analyses were performed using an unpaired two-tailed t-test; ****p < 0.0001; data are shown as mean ± SEM. **(D)** Confocal images of post-mortem human motor cortex from representative controls and sporadic ALS patients immunolabelled with anti-TDP-43 antibody (red), anti-YTHDF3 antibody (green) and anti-MAP2 (violet). DNA was labelled with 4,6-diamidino-2-phenylindole (DAPI) (blue). Images are presented as z-stack projections. Scale bars, 10 µm.

In line with interactomic data and findings from cellular and mouse ALS models, qualitative immunostaining of post-mortem ALS patient tissue also showed altered YTHDF3 immunoreactivity in cortical motor neurons in the presence of TDP-43 aggregates (Figures 7D and S9). When TDP-43 pathology was present, YTHDF3 levels were either reduced or partially colocalized (Figures 7D and S9), with the same pattern observed in TDP-43CTF cells (Figures 6E–6I). These findings shed light on YTHDF3 as an important interactor of TDP-43 in pathology and a possible target for further investigation in ALS/FTD. Furthermore, these results confirm the robustness and quality of our interactomic data.

## Discussion

Our work presents a novel systematic study of the dynamic interactome of TDP-43 at steady state, during oxidative stress, and during recovery, using the APEX2 proximity-labelling method. This method is particularly suitable for studying the interactome at different time points very shortly after induced changes, as labelling of interacting partners occurs rapidly, within one minute. Our data provide a comprehensive view of the interaction environment of TDP-43wt, TDP-43dNLS, and TDP-43CTF before stress, during stress, and throughout the mid- and late recovery phases. These findings provide insights into the processes involving TDP-43 and its variants in physiological and pathological conditions during stress and recovery, and identifies proteins interacting with them during stress and after stress resolution. Given that stress granules can act as seeds for aggregation of TDP-43^44,45^, our in-depth study of the dynamic interactome of TDP-43 and its variants provides a highly useful resource for understanding these processes.

Several studies have addressed the TDP-43 interactome to date^20–22^; however, this is the first study to examine dynamic changes during cell stress and recovery phases after stressor withdrawal. Our APEX2 approach demonstrates that proximity labelling is particularly well suited for studying rapid dynamic processes and identifying transient interactors. This provides a broader understanding of TDP-43 proteinopathies from the prospective of protein-protein interactions and of processes that may be impaired due to ineffective stress response.

Additionally, we identified both known and novel interactors of TDP-43 variants and characterized their dynamic association with these variants by examining their interaction trajectory over time, including whether these interactions became more or less enriched under different conditions. We then classified interactors into clusters according to their interaction trajectory with each TDP-43 variant and observed that our data are consistent with pathological findings. A recent study showed that in FTD-TDP-43 cases, various hnRNPs, including hnRNPA2B1, hnRNPH1, and hnRNPA1, accumulate in the cytoplasm in TDP-43 proteinopathies^46^. This parallels our finding that interaction of TDP-43dNLS with hnRNPA2B1 increased during recovery, whereas its interaction with TDP-43wt decreased, suggesting a greater cytoplasmic abundance of this hnRNP.

Proteinopathies often result from impairment of multiple proteostasis pathways, including the proteasome system^47^, nuclear-cytoplasmic transport^48^, and the ER-Golgi protein secretory pathways^32^. Proteins, the major building blocks of cells, and particularly their interactions, are precisely regulated and are essential for the proper execution of cellular functions and the maintenance of proteostasis^9^. Here, we showed that the TDP-43dNLS and TDP-43CTF variants, whose intracellular behaviour mimics TDP-43 behaviour mimics under pathological conditions, interact with distinct sets of proteins compared to TDP-43wt, both under stress and during recovery. Our results show that both pathology-mimicking forms of TDP-43 interact with proteins involved in RNA processing, homeostasis regulation, and nuclear-cytoplasmic transport, which could lead to impaired stress response, facilitated liquid-liquid phase separation, and aggregation. Moreover, our findings suggest that the cell retains a molecular memory of previous stress within their interactome and mount responses accordingly. These observations support the idea that cellular stress may contribute to the molecular events associated with neurodegeneration.

Having established that stress and recovery reshape the TDP-43 interaction landscape in a variant-specific manner, we next focused on YTHDF3, which displayed a particularly pronounced and biologically relevant interaction pattern in our dataset. Previous studies have shown that TDP-43 physically interacts with m6A writers, including METTL3, as well as with m6A readers such as YTHDF1, YTHDF2, and hnRNPA2B1^26,49^. YTHDF3 is of particular interest, because it functions as m6A reader, serves as a pre-stress granule seed protein^24^, and a component of stress granules with an important role in the cellular stress response. Our data showed that pathology-mimicking forms of TDP-43 interact with YTHDF3 and that this interaction is relevant to disease. Neurons from transgenic mice expressing TDP-43dNLS showed reduced YTHDF3 levels, a finding that we also observed in ALS patients, suggesting that altered YTHDF3 homeostasis may be associated with TDP-43 pathology and highlighting YTHDF3 as a potential therapeutic target.

In summary, our research highlights the complexity and dynamic remodelling of the TDP-43 interactome during stress and recovery. It serves as a detailed resource for future research on the stress-associated dynamic interactome of TDP-43 and its variants. We demonstrated that different cellular states lead to significant rearrangements in the interactomes of TDP-43 variants, which may have pathological consequences. Importantly, this study is among the first to identify the system-wide interactomes of TDP-43wt, TDP-43dNLS and TDP-43CTF, the 25 kDa C-terminal fragment present in toxic aggregates, under steady state, during stress and during stress resolution. Together, these findings provide new insights into the stress-dependent remodelling of TDP-43 interaction networks and their potential contribution to disease-associated TDP-43 dysfunction that may help in developing new therapeutic applications.

## Limitations of the Study

Several limitations of the study should be considered. Interatomic MS studies require high protein yields, so our analysis is limited to human Flp-In HEK293 cells, which allow high protein production. Advances that enable the large-scale production of human neurons or less sample for analysis may help to overcome this technical constraint and facilitate future proteomic studies in more disease-relevant cellular models. In our case, to address this limitation, we validated selected interactor in disease-relevant models in human and mouse brain tissue. To avoid issues associated with stress-related transient transfection, we established inducible stable cell lines.

## STAR Methods

### Key Resource table

| REAGENT or RESOURCE | SOURCE | IDENTIFIER |
|---|---|---|
| Antibodies |  |  |
| Anti-chicken Alexa Fluor 647 | abcam | Cat#<br>ab150171; RRID:AB_2921318 |
| Anti-goat Alexa Fluor 647 | Thermo Fisher Scientific | Cat#<br>A21447; RRID:AB_2535864 |
| anti-MAP2 | Abcam | Cat#ab92434; RRID:AB_2138147 |
| Anti-mouse Alexa Fluor 568 | abcam | Cat#<br>ab175473; RRID:AB_2895153 |
| Anti-mouse Alexa Fluor 647 | Cell Signaling | Cat#<br>4410S |
| Anti-mouse StarBright700 | Bio-Rad | Cat#<br>12004158; RRID:AB_3712167 |
| anti-myc | Proteintech | Cat#60003-2-Ig; RRID:AB_2734122 |
| anti-pTDP-43(ser409/410) | CosmoBio | Cat#CAC-TIP-PTD-M01A; RRID:AB_3251193 |
| Anti-rabbit Alexa Fluor 488 | abcam | Cat#<br>ab150077; RRID:AB_2630356 |
| Anti-rabbit Alexa Fluor 555 | Cell Signaling | Cat#<br>4413S |
| Anti-rabbit Alexa Fluor Plus 488 | Thermo Fisher Scientific | Cat#<br>A32790; RRID:AB_2762833 |
| Anti-rabbit StarBright520 | Bio-Rad | Cat#<br>12005870; RRID:AB_2884949 |
| anti-TDP-43 | Abcam | Cat#ab80608; RRID:AB_1641102 |
| anti-TDP-43 (C-terminal antibody) | Proteintech | Cat#12892-1-AP; RRID:AB_2200505 |
| anti-TDP-43 (N-terminal antibody) | Proteintech | Cat# 10782-2-AP; RRID:AB_615042 |
| anti-YTHDF3 | Proteintech | Cat#25537-1-AP; RRID:AB_2847817 |
| anti-YTHDF3 | abcam | Cat#ab220161; RRID:AB_2868574 |
| Atto 488-Streptavidin | ATTO TEC | Cat#AD 488-61 |
| Streptavidin Protein, HRP | Thermo Fisher Scientific | Cat#21126 |
| Biological samples |  |  |
| Human brain tissue used is in Table S4 | London Neurodegenerative Diseases Brain Bank | N/A |
| Mouse brain tissue used is in Table S4 | Alfieri et al., 2011<sup>70</sup> | N/A |
| Chemicals, peptides, and recombinant proteins |  |  |
| (+)-Sodium-L-ascorbate | Sigma-Aldrich | Cat# A4034-100G; CAS:134-03-2 |
| (±)-6-Hydroxy-2,5,7,8-tetramethylchromane-2-carboxylic acid (Trolox) | Sigma-Aldrich | Cat#238813-25G CAS: 53188-07-1 |
| Biotin Tyramide (Biotin-Phenol) | Sigma-Aldrich | Cat#SML2135-250MG; CAS: 41994-02-9 |
| Doxycycline hyclate | Sigma-Aldrich | Cat# D9891-5G; CAS:24390-14-5 |
| Hydrogen peroxide 30% | Sigma-Aldrich | Cat# H1009-100ML; CAS: 7722-84-1 |
| Sodium arsenite | Sigma-Aldrich | Cat#1.06277.1000; CAS: 7784-46-5 |
| Sodium azide | Sigma-Aldrich | Cat# S2002-25G; CAS: 26628-22-8 |
| Critical commercial assays |  |  |
| Dynabeads MyOne streptavidin T1 | Invitrogen | Cat#65602 |
| Deposited data |  |  |
| Mass spectrometry proteomics data | This paper | PRIDE: PXD078494 |
| Custom code used in this study | This paper | Zenodo: DOI: 10.5281/zenodo.21676737 |
| Experimental models: Cell lines |  |  |
| HEK293 | ATCC | Cat# CRL-1573 |
| SH-SY5Y Flp-In mScarlet-myc-G3BP1 | Motaln et al., 2020<sup>50</sup> | N/A |
| HEK293 Flp-In T-REx-APEX2-myc | This paper | N/A |
| HEK293 Flp-In T-REx-APEX2-myc-TDP-43wt | This paper | N/A |
| HEK293 Flp-In T-REx-APEX2-myc-TDP-43dNLS | This paper | N/A |
| HEK293 Flp-In T-REx-APEX2-myc-TDP-43CTF | This paper | N/A |
| Oligonucleotides |  |  |
| Primers used for cloning are in Table S5 | This paper | N/A |
| Stealth siRNA targeting TDP-43 | Invitrogen | Cat#12935100 |
| Scramble siRNA | Invitrogen | Cat#1299001 |
| Recombinant DNA |  |  |
| pcDNA5/FRT/TO-myc | N/A | N/A |
| pcDNA5/FRT/TO-APEX2-myc | This paper | N/A |
| Flag-APEX2-C | Ken-Ichu Takemaru lab | Addgene plasmid #128146 |
| pcDNA5-BioID2-TDP-43wt | Atwal and Nimac et al. 2025<sup>13</sup> | N/A |
| pcDNA5-BioID-TDP-43dNLS | Atwal and Nimac et al. 2025<sup>13</sup> | N/A |
| pcDNA5-mScarlet-myc-TDP-43wt | Atwal and Nimac et al. 2025<sup>13</sup> | N/A |
| pcDNA5-mScarlet-myc-TDP-43dNLS | Atwal and Nimac et al. 2025<sup>13</sup> | N/A |
| pcDNA5-mScarlet-myc-TDP-43CTF | This paper | N/A |
| pT7-EGFP-C1-HsYTHDF3_AB | Elisa Izaurralde lab | Addgene #148313 |
| pcDNA5-myc-TDP-43wt | This paper | N/A |
| pcDNA5-myc-TDP-43dNLS | This paper | N/A |
| pcDNA5-myc-TDP-43CTF | This paper | N/A |
| Software and algorithms |  |  |
| MaxQuant | Cox et al. 2008<sup>54</sup> | https://maxquant.org |
| Andromeda search engine | Cox et al. 2011<sup>56</sup> | N/A |
| Perseus | Tyanova et al. 2016<sup>58</sup> | http://maxquant.org/perseus |
| Fiji (ImageJ) software | Schindelin et al. 2012<sup>69</sup> | https://imagej.net/software/fiji/ |
| Cytoscape | Kohl et al. 2011<sup>62</sup> | Version 3.10.4; https://cytoscape.org/ |
| Image Lab Software | Bio-Rad Laboratories | https://www.bio-rad.com/en-si/product/image-lab-software?ID=KRE6P5E8Z |
| AlphaFold3 | Abramson et al. 2024<sup>52</sup> | https://alphafoldserver.com/ |
| R | R Foundation for Statistical Computing | https://www.r-project.org/?utm_source=chatgpt.com, Version 4.4.1 |
| GraphPad Prism | GraphPad Software | https://www.graphpad.com/features |
| CellSense Dimension | Evident Scientific (formerly Olympus) | https://evidentscientific.com/en/products/software/cellsens |

## Resource Availability

### Lead contact

All information and other reasonable requests for data and reagents will be made available by contacting the Lead Contact, Boris Rogelj. Materials used in this study will be available upon request from the Lead Contact, under a standard MTA.

### Data availability

Proteomics raw data generated in this study are available via PRoteomics IDEntifications database (PRIDE repository), under accession number PXD078494.

#### Code availability

Source code used for analysis are accessed from GitHub (DOI: 10.5281/zenodo.21676737).

### Experimental Model and Subject details

#### Mammalian cell cultures

HEK293 cell lines were obtained from the American Type Culture Collection (ATCC). Cells tested negative for mycoplasma contamination, confirmed by Eurofins Genomics and were cultured at 37°C in a humidified atmosphere with 5% CO_2_. Cells were grown in Dulbecco’s modified Eagle medium with high glucose, as GlutaMAX supplement (DMEM, Gibco, 10566016), supplemented with 10% fetal bovine serum (FBS, Gibco, 10270-106) and 100 U/mL penicillin-streptomycin (Gibco, 15140122). All established HEK293 Flp-In T-REx cell lines were grown under the same conditions as HEK293 cells, except that tetracycline-free 10% fetal bovine serum (FBS Tet Free, Euroclone, ECS0182L) was used instead of standard FBS, and cells were cultured in 200 µg/mL hygromycin B (InvivoGen, ant-hg-1) and 10 µg/mL blasticidin S (Sigma-Aldrich, SBR00022-1ML). SH-SY5Y Flp-In mScarlet-myc-G3BP1^50^ cells were cultured under the same conditions as HEK293 cells, except for the culture medium. These cells were grown in Dulbecco’s modified Eagle medium F12 (DMEM F12, Gibco, 11320033), supplemented with that tetracycline-free 10% fetal bovine serum (FBS Tet Free, Euroclone, ECS0182L) and 100 U/mL penicillin-streptomycin (Gibco, 15140122).

#### Mouse brain tissue

TDP-43dNLS bigenic mice (n = 4) and corresponding control animals (n = 4) were used in this study. Both male and female mice were included. Detailed information on the experimental animals, is provided in Table S4. All animal experiments were approved by the National Animal care and Use Committee (CICUAL) of the School of Medicine, University of Buenos Aires (UBA) (RESCD-2023-2358-E-UBA-DCT#FMED).

#### Human brain tissue

Human post-mortem motor cortex samples were sourced from the London Neurodegenerative Diseases Brain Bank (King’s College London, UK), which operates with consent for autopsy, neuropathological assessment, and research for all donors. Post-mortem motor cortex tissue from individuals with ALS (n = 6) and neurologically healthy controls (n = 6) was analysed. Clinical and demographic characteristics of individual cases, including age, sex, diagnosis, post-mortem interval, and neuropathological information, are provided in Table S4. All studies were conducted with the ethical approval from the tissue bank (23/WA/0124).

## Method details

### Generation of plasmids

Primer sequences (Table S3) were used for Gibson Assembly (GenScript, L00701) into pcDNA5/FRT/TO-myc or pcDNA5/FRT/TO-APEX2-myc. The gene sequence of APEX2 was amplified by PCR from the Flag-APEX2-C cloning vector. The Flag-APEX2-C cloning vector was a gift from Ken-Ichu Takemaru (Addgene plasmid #128146). The restriction enzymes used for APEX2 insertion into pcDNA5/FRT/TO-myc were BamHI and EcoRV (Thermo Scientific), while NotI and PspOMI (Thermo Scientific) were used to insert the TDP-43 variant into pcDNA5/FRT/TO-APEX2-myc. The gene sequences of the TDP-43 variants were amplified by PCR from pcDNA5-BioID2-TDP-43wt or pcDNA5-BioID-TDP-43dNLS. To generate myc-tagged TDP-43 constructs, TDP-43 variants were amplified by PCR from pcDNA5-mScarlet-myc-TDP-43wt, pcDNA5-mScarlet-myc-TDP-43dNLS or pcDNA5-mScarlet-myc-TDP-43CTF (Table S3). The newly constructed plasmids were verified by Sanger sequencing (Eurofins).

### Stable cell line generation

Stable cell lines were established using the HEK293 Flp-In T-REx system (Invitrogen) following the manufacturer’s protocol. HEK293 Flp-In T-REx cells were co-transfected with either pcDNA5/FRT/TO-APEX2-myc, pcDNA5/FRT/TO-APEX2-myc-TDP-43wt, pcDNA5/FRT/TO-APEX2-myc-TDP-43dNLS, or pcDNA5/FRT/TO-APEX2-myc-TDP-43CTF, and pOG44 vectors at a 1:9 ratio using PolyJet In Vitro DNA Transfection Reagent (SignaGen Laboratories, SL100688), followed by hygromycin B (InvivoGen, ant-hg-1) selection and single-cell selection. Stable cell lines HEK293 Flp-In T-REx-APEX2-myc, HEK293 Flp-In T-REx-APEX2-myc-TDP-43wt, HEK293 Flp-In T-REx-APEX2-myc-TDP-43dNLS, and HEK293 Flp-In T-REx-APEX2-myc-TDP-43CTF were generated. Expression of APEX2-myc, APEX2-myc-TDP-43wt, APEX2-myc-TDP-43dNLS and APEX2-myc-TDP-43CTF was induced with 1 µg/mL doxycycline (Sigma-Aldrich, D9891-5G).

### Western blot analysis

Eluates prepared after pull-down assay in 2×SDS loading buffer with 200 mM DTT, incubated at 95°C for 10 min were resolved on 12% stain-free gels (Bio-Rad) at 120 V, followed by a semi-dry transfer onto a nitrocellulose membrane (GE Healthcare, 10600008) at 1.3 A per membrane for 10 min. The nitrocellulose membrane was blocked in blocking solution (5% skimmed milk in TBS with 0.05% Tween-20 (TBST), or 5% BSA in TBST for biotinylated proteins). Incubations with primary and secondary antibodies (Table S5) were carried out as suggested by the manufacturer. For signal detection, Clarity Western ECL Substrate (Bio-Rad) was used. Images were acquired using the GelDoc System (Bio-Rad) and ImageLab (Bio-Rad) software was applied for densitometric analysis.

### Immunostaining

Cells were grown on coverslips treated with poly-L-lysine in 24-well plates at a density of 1 × 10^5^ cells per well. After twenty-four hours of fusion protein expression induction or twenty-four hours following transient transfection with Lipofectamin 3000 (Invitrogen, L3000015), cells were washed with DPBS (Gibco, 14190-169) and fixed with 4% paraformaldehyde (PFA) solution diluted in DPBS for 15 min. Blocking and permeabilization were performed in blocking solution containing 5% horse serum (HS, Gibco, 26050-088) and 0.1% Triton X-100 in DPBS for 30 min at room temperature. Cells were then incubated with primary antibody solution diluted in blocking solution according to the manufacturer’s protocol for 60 min at room temperature. After washing with DPBS, coverslips were incubated with secondary antibody solution diluted in blocking solution according to the manufacturer’s protocol for 60 min at room temperature. The list of primary and secondary antibodies is provided in Table S5. Nuclear counterstaining was achieved by incubation with DAPI for 7 min. Cells were mounted in ProLong Gold mounting medium (Invitrogen, P36930). Imaging was performed using a Zeiss LSM 710 inverted confocal laser scanning microscope with a Plan-Apochromat 63×/1.4 NA M27 oil immersion objective and immersion oil (Carl Zeiss, 4Y00-R0DY-1007-3VF3). DAPI, Alexa Fluor 488, Atto 488, Alexa Fluor 555, and Alexa Fluor 647 were excited using wavelengths of 405, 488, 543 and 647 nm, respectively. The zoom factor was adjusted between 1 and 3.2×, and images were acquired at 1024 × 1024 pixels along both axes with ZEN 2011 image software (Carl Zeiss). ImageJ was used for image export, and for fluorescence intensity profiles (Table S6).

### APEX2 method

The APEX2 method was carried out as previously described by Marmor-Kollet et al. (16). In brief, HEK293 Flp-In T-REx cells expressing TDP-43 variants fused to APEX2 or APEX2-myc were grown in 10-cm dishes in triplicate for each condition and construct, totalling 48 dishes. When the cells reached 70−80% confluence, expression of APEX2-myc, APEX2-myc-TDP-43wt, APEX2-myc-TDP-43dNLS, and APEX2-myc-TDP-43CTF was induced. Twenty-four hours later, all cells except those used to study the steady state interactome were exposed to 50 µM sodium arsenite (SA, Sigma-Aldrich, 1.06277.1000) for 90 min at 37 °C in a humidified atmosphere with 5% CO_2_. Ninety minutes after SA exposure, the medium was changed for cells in which the interactome was studied 4 h and 24 h after SA removal corresponding to 50% stress granule burden and full recovery, respectively. Sixty minutes after SA exposure, 500 µM biotin-phenol was added to the cells used to study the interactome under stress conditions and steady state for 30 min at 37 °C in a humidified atmosphere with 5% CO_2_. Interaction labelling was performed with 100 mM H_2_O_2_ for 1 min at room temperature. The reaction was stopped by rapid medium aspiration and addition of cold quencher solution (5 mM Trolox, 10 mM sodium ascorbate, 10 mM sodium azide in DPBS). Cells were washed three times, scraped off, and centrifuged at 300 g at 4 °C for 10 min. The pellets were freshly frozen in liquid nitrogen to ensure identical conditions for all time points at which cells were harvested. The procedure was repeated 4 h and 24 h after stressor removal to label interaction partners at these time points.

### Pull down assay

Freshly frozen pellets were thawed on ice and lysed in urea lysis buffer (8 M urea, 50 mM Tris, pH 7.4, 5 mM Trolox, 10 mM sodium ascorbate, 10 mM sodium azide, and 1 mM DTT) supplemented with protease inhibitors (Roche, 05892791001). After resuspension, cells were left on ice for 30 min. Immediately before sonication 20% Triton X-100 was added to the lysates. Lysates were sonicated at medium intensity for 20 sec, followed by a 20 sec pause, and this cycle was repeated three times. Lysates were then cleared by centrifugation at 20,000 × g for 30 min at 4 °C. The supernatant was transferred to fresh low-binding tubes. Magnetic Dynabeads MyOne Streptavidin T1 beads (Invitrogen, 65602) were equilibrated in 50 mM Tris, pH 7.4, three times. Whole lysates were incubated with 100 µL magnetic Dynabeads MyOne Streptavidin T1 beads per sample overnight at 4 °C using end-over-end rotation. The following day, magnetic Dynabeads MyOne Streptavidin T1 beads were collected on a magnetic rack and washed in a series of buffers (Table S5) to remove nonspecific binders. After the final wash, beads were resuspended in 50 mM ammonium bicarbonate and flash frozen before mass spectrometry analysis, whereas when proceeded with western blot analysis for validation the beads were resuspended in 1× Laemmli buffer.

### Subcellular fractionation

Subcellular fractionation was performed as previously described by Prpar Mihevc et al.^51^. Briefly, cells were cultured in six-well plates, and on the day before harvesting, protein expression was induced with doxycycline. After 24 h, cells were lysed in ice-cold lysis buffer (50 mM Tris, pH 7.4, 10 mM NaCl, 0.5% IGEPAL CA-630 (Sigma-Aldrich, 18896-100mL), 0.25% Triton X-100, protease inhibitors and phosphatase inhibitors (Roche)) and centrifuged at 3,000 × g for 5 min at 4 °C. The resulting supernatant was transferred to fresh tubes and re-centrifugated at 16,100 × g for 10 min at 4 °C. The cleared supernatant was the cytoplasmic fraction. The pellet remaining after the first centrifugation was washed three times in cell lysis buffer, then resuspended in in urea lysis buffer, sonicated, and centrifuged again at 16,100 × g for 10 min at 4 °C. This was the nuclear fraction. A pull-down assay was performed to isolate interaction partners of TDP-43. Eluates from the pull-down assay were resolved by SDS-PAGE, followed by western blot analysis.

### Knockdown assay

HEK293 cells were 70% confluent on the day of siRNA transfection with Lipofectamine 3000 (Invitrogen, L3000001). Transfection was performed according to the manufacturer’s protocol. Stealth siRNA targeting TDP-43 and Scramble siRNA were obtained from Invitrogen (12935100 and 1299001, respectively). Cells were collected 48 h post-transfection for WB analysis and knockdown efficiency. The experiment was performed in three independent biological replicates, each with three technical replicates (Table S6). Technical replicates were averaged before statistical analysis. Knockdown and siScramble groups were compared using a paired two-tailed t-test.

### AlphaFold interaction prediction

The predicted structure of the protein complex was generated using AlphaFold 3^52^ via AlphaFold Server using default settings. Full-length protein sequence was submitted as separate chains, the top-ranked model was selected based on the server confidence ranking and visualized in UCSF ChimeraX^53^.

### Silver staining

After the pull-down assay and before mass spectrometry analysis, we took 5% of the eluate to assess the quality of the sample for mass spectrometry using silver staining. Samples were diluted in 2×SDS loading buffer with 200 mM DTT, incubated at 95°C for 10 min and resolved on 12% polyacrylamide precast gels (Invitrogen) at 120 V, followed by two-hours of gel fixation in 45% methanol and 5% glacial acetic acid diluted in ultrapure water. The sensitization step was performed by incubating gel in Na_2_S_2_O_3_ for 1 min at room temperature, followed by three sequential rinses in ultrapure water. Next, incubation in AgNO_3_ and 0.06% formaldehyde in the dark at room temperature was performed. After a further three sequential rinses in ultrapure water, development was carried out with developing solution (Na_2_CO_3_, formaldehyde, methanol, and glacial acetic acid). When bands of biotinylated proteins appeared, the reaction was stopped with 10% glacial acetic acid diluted in ultrapure water. Images were acquired using the GelDoc System (Bio-Rad), and ImageLab (Bio-Rad) software was used for densitometric analysis.

### MS sample preparation

25 µL of enriched sample coupled to magnetic beads (Dynabeads MyOne streptavidin T1) was digested in solution by adding 50 mM ammonium bicarbonate and 0.3 µg of Trypsin (trypsin:protein ratio, 1:25). Digestion was carried out overnight in a thermocycler at 37°C and gentle shaking. Digestion was stopped by acidification by adding 10% trifluoro acetic acid (final concentration 0.5%). Samples were recovered on a magnetic rack and the supernatant was transferred to a new Eppendorf tube. Peptides were dried in a vacuum centrifuge and dissolved in 20 µL 0.1% TFA.

### Mass spectrometric measurements

Liquid chromatography (LC) was performed using the Vanquish Neo UHPLC system (Thermo Fisher Scientific, Bremen, Germany). Approximately 200 ng of peptides were loaded onto a precolumn (PepMap Neo C18 Trap Cartridge, 300 μm × 0.5 cm, particle size 5 μm) and then separated on an analytical column (DNV PepMapTM Neo, 75 μm × 150 mm, C18, particle size 2 μm, pore size 100 Å). Peptides were separated at a flow rate of 400 nl/min using a solvent gradient from 1% B to 21% B (B: 84 % acetonitrile, 0.1% formic acid) over 70 min and a subsequent increase to 40% B for 20 min. The column was rinsed with 99 % B for 5 min. Subsequently, the column was adjusted back to 5% solution B. The LC system was online coupled an Orbitrap Fusion Lumos Tribrid mass spectrometer (Thermo Fisher Scientific, Bremen, Germany). Samples were measured in data dependent acquisition. MS1 scans were carried out at a resolution of 60,000 a normalized AGC target of 250% an RF lens of 45% and an injection time of 50 ms. MS1 scans were filtered for a minimum intensity of 1 × 10^5^ and a maximum intensity of 1 × 10^20^ and charge states between 2-7. The dynamic exclusion was set to 80 s. Subsequently the Top 15 MS1 scans were selected for fragmentation. MS2 scans were carried out with a resolution of 30,000 a normalized AGC target of 100% and a maximum injection time of 120 ms. Fragments were generated by HCD with a normalized collision energy of 32%.

### MS data analysis and data screening

Data analysis was carried out in MaxQuant (version 2.4.9.0)^54^. Resulting peak lists were searched against the human SwissProt FASTA (Uniprot, accessed 15. 12. 2023), containing additionally all known TDP43 variants and a common contaminants database provided in the Andromeda search engine^56^. Oxidation of methionine was set as variable modification due to sample preparation. Trypsin was set as digestion enzyme and a maximum number of 2 missed cleavages were tolerated in the search. The false discovery rate (FDR) was set to 1% for peptides and proteins. Peptide identification was carried out with an initial allowed precursor mass deviation up to 5 parts per million (ppm) and an allowed fragment mass deviation of 20 ppm. Match between runs was disabled. The resulting proteingroups.txt output was imported into Perseus (version 2.0.10.0)^58^. Contaminants and decoys were removed. Statistical analyses and graphing for Figures 2-4A, 4B, 4E and Figure 5, as well as Figures S3-S5D and S5G and as Figure S6 were conducted using R version 4.4.1^55^ within RStudio.

The protein intensities were log2-transformed and normalized using the median, quantile, and loess normalization approaches^57^. After assessment of the boxplots, PCA-plots, and MA-plots, loess normalization was chosen for the further analyses. The normalized intensities were calculated once on the whole dataset and once for each experimental group (APEX, CTF, dNLS, and WT) separately.

The protein intensities that were normalized over the whole data set were used to calculate ANOVAs comparing the different experimental groups at one specific timepoint, assuming unequal variances (Welch-ANOVA) using the ProtStatsWF R package, version 0.0.0.9000^59^. The ANOVA p-values were corrected for multiple testing using the Benjamini-Hochberg procedure. Post-hoc tests to compare the time points in a pairwise manner were also conducted. Proteins with a corrected ANOVA p-value < 0.05, a corresponding posthoc p-value < 0.05, and a fold change of >2 or < 0.5 were considered as significant.

The intensities that were normalized per group were used to compare the different timepoints within one experimental group, again using Welch ANOVAs. Additionally, agglomerative hierarchical clustering with complete linkage and distance based on Pearson-correlation was applied and visualized using heatmaps (ComplexHeatmap R package, version 2.18.0^60^). For trajectory presentation, we defined clusters in the Orange Data Mining tool^61^ by cutting the hierarchical clustering dendrogram at a distance threshold selected using the elbow rule. Selected clusters were visualized in R using ggplot2 package^60^.

### On/Off analysis

Proteins were classified into phase-specific interaction patterns based on detection across unstressed conditions, under oxidative stress, early recovery and late recovery. A protein was considered detected in a condition if it was identified in at least 2 out of 3 replicates. Detection was then converted into a binary presence/absence pattern across the time course. The 1111 binary pattern was classified as constitutive; stress-associated patterns included proteins detected during the stress phase (0100, 0110, 0111, 0101); the 0010 pattern indicated recovery-specific proteins; sustained recovery-associated proteins followed the pattern 0011 pattern; late recovery-specific proteins were 0001; and the 1000, 1100, 1001, 1011 and 1101 patterns were associated with stress-sensitive/lost proteins. Proteins with the 0000 pattern were excluded from the On/Off analysis, as they were not detected at any time point. The output data table is provided in Table S1.

### Network visualization

The interaction networks of different clusters that follow the same interaction trajectory with TDP-43 variant were visualized using Cytoscape version 3.10.4 (https://cytoscape.org/)^62^.

### GO analysis and GO-based descriptive annotation analysis

GO analysis was performed using g: Profiler in R (package gprofiler2, version 2.0.2.4)^63^, with q-values (adjusted p-values using the Benjamini-Hochberg false discovery rate method) < 0.05. Only significantly enriched terms passing the FDR threshold were retained. GO analysis included cellular compartment, molecular function, biological process, Reactome, and KEGG pathways using TDP-43 variant-specific interactors (compared to TDP-43 variant or control at different time points) with absolute log2FC ≥ 2 and posthoc p-value < 0.05. Complete enrichment results for each comparison are provided in Table S2. For graphical representation, significantly enriched terms were visualized as dot plots in ggplot2 R package^64^, with the side representing gene ratio, dot colour representing q-value, and the x-axis representing the number of overlapping genes.

Candidate proteins from the late recovery-associated and persistent post-stress sets were further analysed using an automated descriptive GO-based annotation workflow. This was a rule-based profiling approach to summarise functional and compartmental features. Candidate proteins were mapped to Entrez identifiers using the org.Hs.eg.db package^65^, and GO annotations retrieved with AnnotationDbi^66^ and GO.db package^67^. GO terms were collapsed per protein and separated by ontology into biological process, molecular function and cellular component terms. Functional categories were assigned by matching keywords across all GO term text, whereas component categories were assigned by matching keywords against cellular component GO terms. Detailed results, including all GO terms, are provided in Table S3.

### Temporal classification of stress and recovery associated protein abundance changes

Protein-level temporal responses were analysed separately for TDP-43wt, TDP-43dNLS and TDP-43CTF variants using triplicate z-score abundance values – for unstressed (0 h), 100% SG (1.5 h), 50% SG (4 h) and recovery (24h) conditions. For each protein, mean z-scores were calculated at each time point from valid replicate values, and changes were defined relative to steady state as Δ1.5 h, Δ4 h and Δ24 h. Proteins were classified as showing persistent post-stress changes if absolute (Δ) values met thresholds of ≥ 1.1, ≥ 0.9 and ≥ 1.0 at 1.5 h, 4 h and 24 h, respectively. These proteins also retained 0.6-1.6-fold of the 1.5 h change. With this we excluded transient responses that strongly declined by recovery and delayed responses that were weak at the beginning but stronger at recovery. Moreover, persistent post-stress changes were required to show same direction of regulation from stress onwards. Proteins were classified as late recovery-associated if early changes were limited (IΔ1.5hI ≤ 0.5 and IΔ4hI ≤ 0.7) but the recovery change was stronger (IΔ24hI ≥ 1.3 and IΔ4hI ≥ 0.7 greater than the maximum early change). The criteria for late recovery-associated proteins excluded those with strong early responses and retained those whose responses appeared during recovery.

Proteins were ranked using weighted change scores that prioritised sustained or late-emerging changes and penalised inconsistent direction or large early responses, respectively. Temporal changes were calculated as follows:

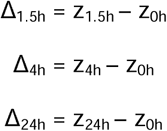

Thresholds were selected empirically to identify robust and interpretable temporal response profiles from normalized protein abundance z-score values. Changes in mean z-score values were calculated against the steady state for each variant. Proteins were assigned to response classes based on the magnitude, timing and direction of these changes. The aim of this approach was to define biologically relevant temporal categories, including persistent post-stress changes and late recovery-associated changes. Cut-offs near one z-score unit were used to focus on changes exceeding minor baseline variation.

For Venn diagrams, an additional display-strict filter was applied after the main temporal classification. Proteins were required to be present at all time points with at least 2 of 3 valid replicates per time point, for a variant to be included in Venn display sets. Persistent post-stress proteins were included in the Venn diagram only if they met the main persistent classification and showed an absolute change of at least 1.0 at each post stress time point. Late recovery associated interactors were included in the Venn diagram with stricter thresholds than in the main classification. This required an absolute recovery change of at least 1.4 and a recovery change exceeding the maximum earlier change by at least 0.8 z-score units.

### Live imaging of YTHDF3 and TDP-43dNLS

HEK293 cells were 70% confluent on the day of co-transfection with pT7-EGFP-C1-HsYTHDF3_AB and pcDNA5-mKO2-TDP-43dNLS^68^ using Lipofectamine 3000 (Invitrogen, L3000001). pT7-EGFP-C1-HsYTHDF3_AB was a gift from Elisa Izaurralde (Addgene #148313). The cells were live-imaged at 37°C and 5% CO_2_, 24 h post-transfection, using Olympus IXplore83 microscope equipped with CSU-W1 spinning disc and an Okolab H301 K-frame live imaging chamber. Selected regions of interest were imaged using the 60×/1.4 NA Oil immersion lens and the CellSens Dimension software platform (Olympus) at 15 min intervals from the addition of 50 μM sodium arsenite until the final timepoint at 90 min. Untreated cells were used as a control. Fluorescence intensity profiles (Table S6) of EGFP-YTHDF3 and mKO2-TDP-43dNLS were generated using Fiji software^69^.

### Immunohistochemistry of human brain tissue

Human post-mortem motor cortex samples were sourced from the London Neurodegenerative Diseases Brain Bank (King’s College London, UK), which operates with consent for autopsy, neuropathological assessment, and research for all donors. All studies were conducted with the ethical approval from the tissue bank (23/WA/0124). Sections (7 µM) were prepared from paraffin-embedded tissue blocks (10% formalin-fixed) and mounted on slides for subsequent staining. Sections underwent sequential de-waxing steps in xylene tanks (2× for 2 min), 99% Industrial Methylated Spirit (IMS, 2× for 2 min) and finally 95% IMS (1× for 2 min) before being rinsed under running water and then moved to distilled water. To unmask antigen epitopes, tissue sections underwent a microwave enhanced antigen retrieval protocol with citrate buffer (0.01M, pH 6.0) by microwaving at high power for 6 min, followed by two 8 min cycles at low power. After cooling, slides were rinsed with phosphate-buffered saline (PBS) before immunostaining. This was followed by blocking with 10% Normal Goat Serum (ab7481, Abcam) in PBS for approximately 45 minutes. Tissue sections were then incubated with primary and secondary antibodies (Table S5) according to the manufacturer’s protocol (with a 2×, 5 min each washing step in between primaries and secondaries in PBS). After another 2×, 5 min washing step in PBS, sections were incubated in quenching buffer (Biotium, 23014) for 8 min at room temperature followed by a 2× wash in PBS for 5 min. To counterstain nuclei, sections were incubated with DAPI for 10 min. Finally, the sections were directly mounted with EverBrite Hardset Mounting Medium (Biotium, 23003). Images were captured on a Nikon Upright Ni-E with A1R confocal optics, through a 60x oil 1.4NA objective. The clinical details of the cases used in this study are provided in Table S4.

### Mouse brain tissue collection

All animals were deeply anesthetized by intraperitoneal injection of 5% chloral hydrate (1 ml per 30 g body weight) and transcardially perfused with ice-cold PBS (0.1 M, pH 7.4) containing 10 U/mL heparin. Brains were rapidly dissected, removed, and fixed overnight in 4% paraformaldehyde (PFA). After fixation, mouse brain tissue was cryoprotected in 10% or 30% sucrose in PBS. Using a sliding freezing microtome (SM 2010R; Leica), fixed frozen brain hemispheres were cryosectioned at 50 µm. All animal experiments were approved by the National Animal care and Use Committee (CICUAL) of the School of Medicine, University of Buenos Aires (UBA) (RESCD-2023-2358-E-UBA-DCT#FMED).

### Immunohistochemistry of mouse brain tissue

Mouse brain tissue was used for immunofluorescence studies as previously described in^70^. Briefly, free-floating mouse brain sections were washed in DPBS (Gibco, 14190-169) and permeabilized in 1% Triton X-100 in DPBS for 1 h. Blocking was performed for 1 h at room temperature in blocking solution (5% horse serum (HS, Gibco, 26050-088), 0.3% Triton X-100 in DPBS). Mouse brain sections were then incubated with primary antibodies (Table S5) diluted in 3% HS, 0.3% Triton X-100 in DPBS for 3 h at room temperature, followed by overnight incubation at 4°C. After 3× washes in 0.04% Triton X-100 in DPBS, slices were incubated with secondary antibodies (Table S5) diluted in 3% HS, 0.3% Triton X-100 in DPBS for 4 h at room temperature, followed by washing in 0.04% Triton X-100 in DPBS. For nuclear counterstaining Hoechst 33342 was used (30 min at room temperature). Mouse brain sections were mounted in EverBrite Hardset Mounting Medium (Biotium, 23003) on gelatin-coated slides (Epredia, J1800AMNZ). Images were captured with an Olympus IXplore IX83 Microscope using the 60×/1.4 NA Oil immersion lens and the CellSens Dimension software platform (Olympus). For z-stacks, settings were kept constant across individual z-planes. Fiji software^69^ was used to linearly enhance the contrast and brightness of individual channels and for image export. Images were processed using ImageJ, with YTHDF3 intensity, neuronal, nuclear and cytoplasmic area, and nuclear-cytoplasmic ratios generated from pixel intensities, which were used as a quantitative readout for protein expression (Table S6). Brain tissue from four mice (Table S4) per group was analysed. Statistical analyses were performed using an unpaired two-tailed t-test.

### Statistics and reproducibility

Unless specified, quantitative data was analysed using GraphPad Prism 11.0.2 software.

## Acknowledgments

This research was supported by the FEBS Summer Fellowship to JN. We are grateful to Dr. George Chennell of the Wohl Cellular Imaging Centre at King’s College London (including UpRight microscope) for technical support. Human tissue samples were provided by the London Neurodegenerative Diseases Brain Bank, King’s College London which receives partial funding through the Brains for Dementia Research Project (jointly funded by the Alzheimer’s Society and Alzheimer’s Research UK). Some graphics were created in BioRender.

## Author contributions

Conceptualization: JN, BR; Methodology: JN, SL, LB, KS, HM, KP, AMK, UC, KRF, FV, LM, LMI, ME, KM; Validation: JN, SL, LB; Formal analysis: JN, SL, LB, BE, KS, KP, AMK, FV, LM, LMI; Resources: LMI, BNS, ME, KM, CT, JU and BR. Writing – original draft: JN; Writing – review and editing: JN, SL, LB, BE, KS, HM, AMK, KP, UC, KRF, BNS, VZ, FV, LM, LMI, ME, KM, CT, JU and BR. Supervision: BR, CT, JU, LMI, BNS, ME, KM; Project administration: BR; Funding acquisition: LMI, BNS, ME, KM, CT, JU, BR.

## Ethic statement

The human post-mortem motor cortex samples were provided by the London Neurodegenerative Disease Brain Bank (King’s College London, UK). Consent for autopsy, neuropathological assessment and research was obtained for all donors and all studies were carried out under ethical approval of the tissue bank (23/WA/0124 REC 3 for Wales). All animal experiments were approved by the National Animal care and Use Committee (CICUAL) of the School of Medicine, University of Buenos Aires (UBA) (RESCD-2023-2358-E-UBA-DCT#FMED).

