## supplementary material for "A temporal atlas of TDP-43 interactions reveals persistent network remodelling beyond stress granule clearance"

**Figures: 7**

**Supplementary figures: 9**

**Supplementary tables: 5**

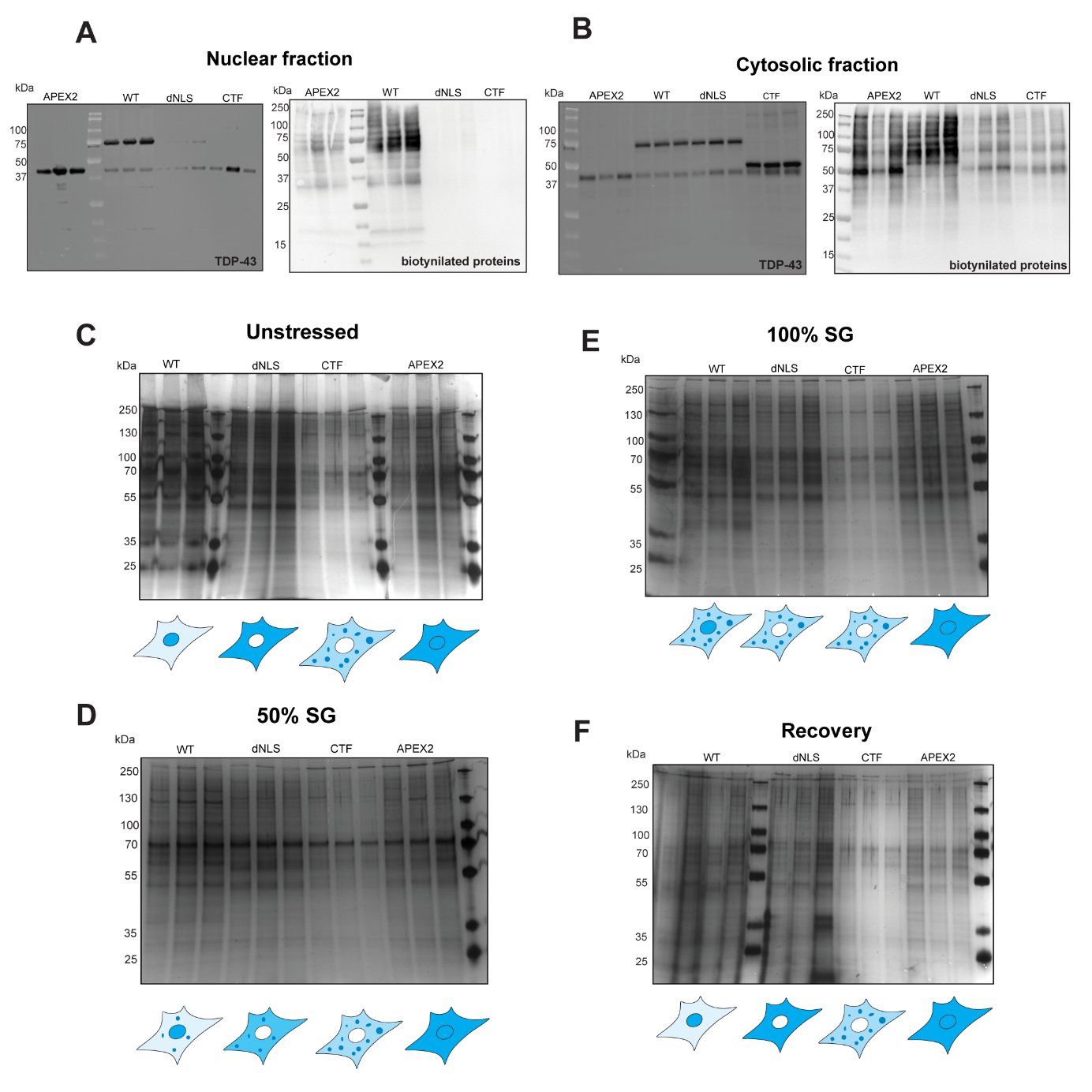

**Figure S1: Fractionation controls and silver staining results support construct-specific proteomic differences. (A-B)** Using fractionation followed by pull-down assay, we demonstrated the quality of established stable cell lines. **(A)** In the nuclear fraction, we detected TDP-43 in APEX2 and TDP-43wt; in the same samples we also detected biotinylated proteins. **(B)** In the cytosolic fraction, we detected TDP-43 and biotinylated proteins in all samples. **(C-F)** Silver staining of biotinylated proteins isolated in the pull-down assay**.** The experiment was performed in three technical replicates. **(C)** Unstressed cells, **(D)** cells with 100% stress granule burden, **(E)** cells with 50% stress granule burden, and **(F)** recovery.

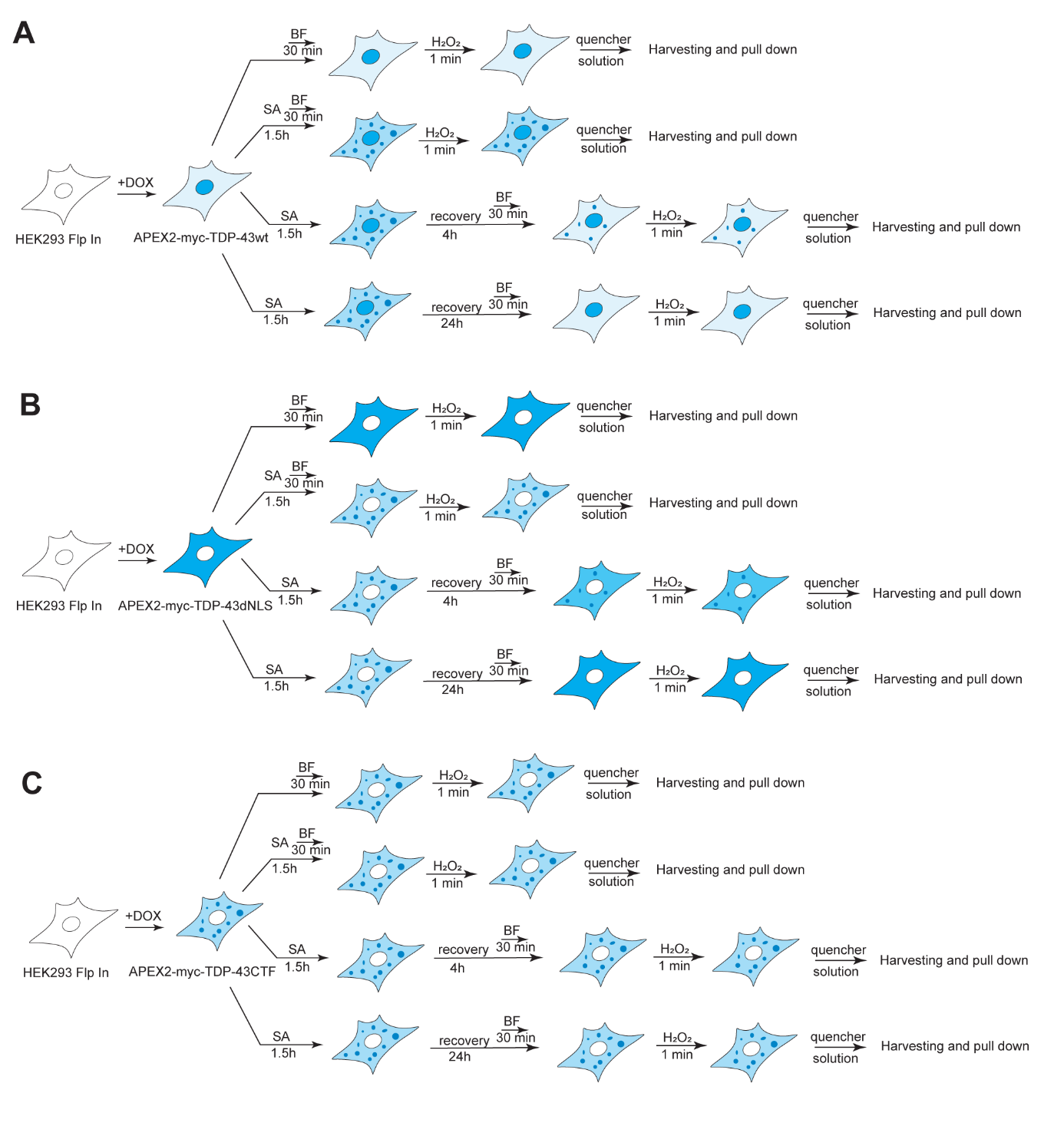

**Figure S2: Experimental workflow of APEX2 from steady state through stress to recovery.** Overview and expected behaviour of **(A)** TDP-43wt**, (B)** TDP­-43dNLS**,** and **(C)** TDP-43CTF**.**

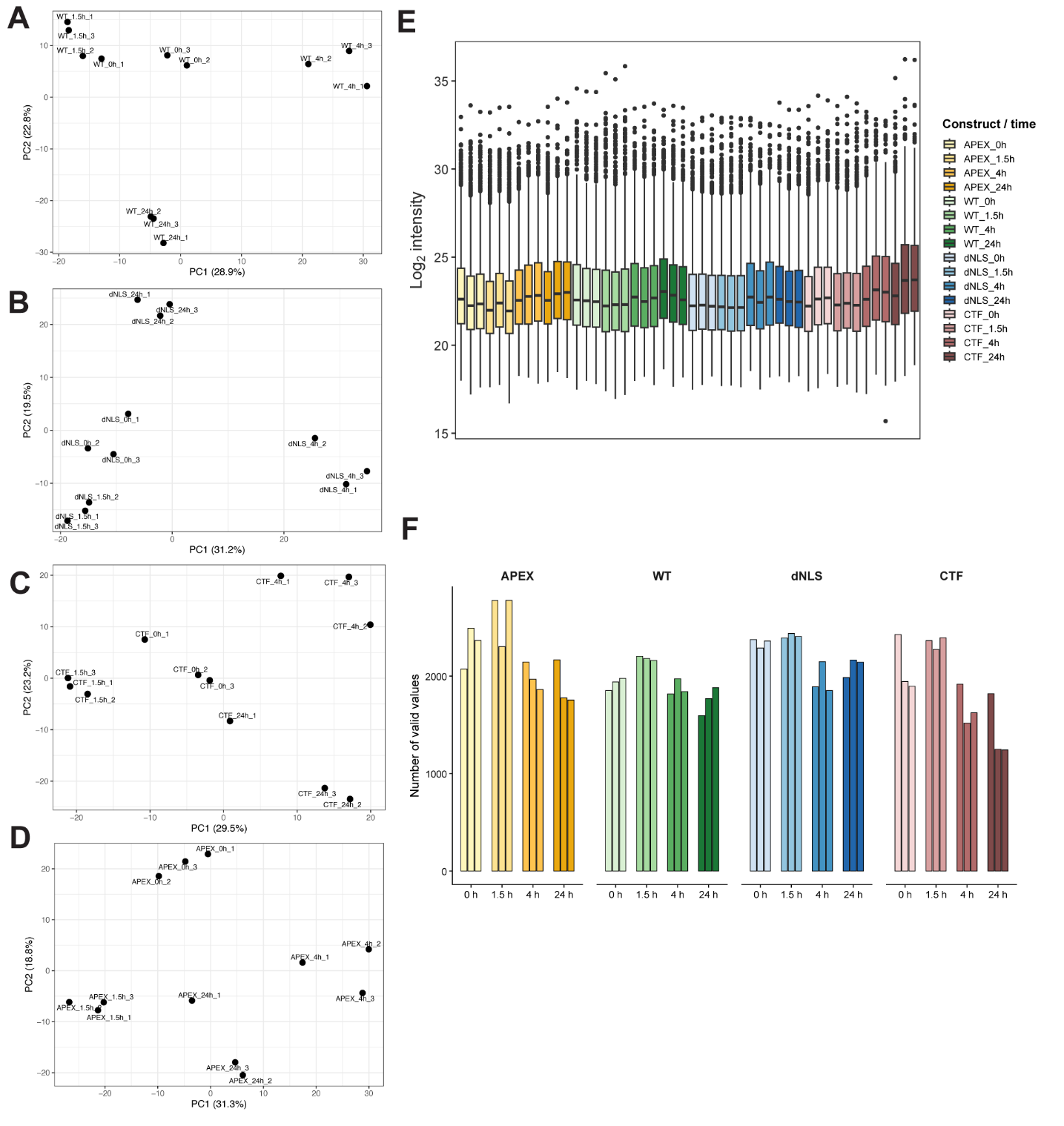

**Figure S3: Principal component analysis (PCA) and proteomic data quality metrics. (A-D)** PCA analysis of proteomic samples showing separation according to time point. **(E-F)** quality control plots show **(E)** log2 intensity distributions and **(F)** the number of valid values across samples.
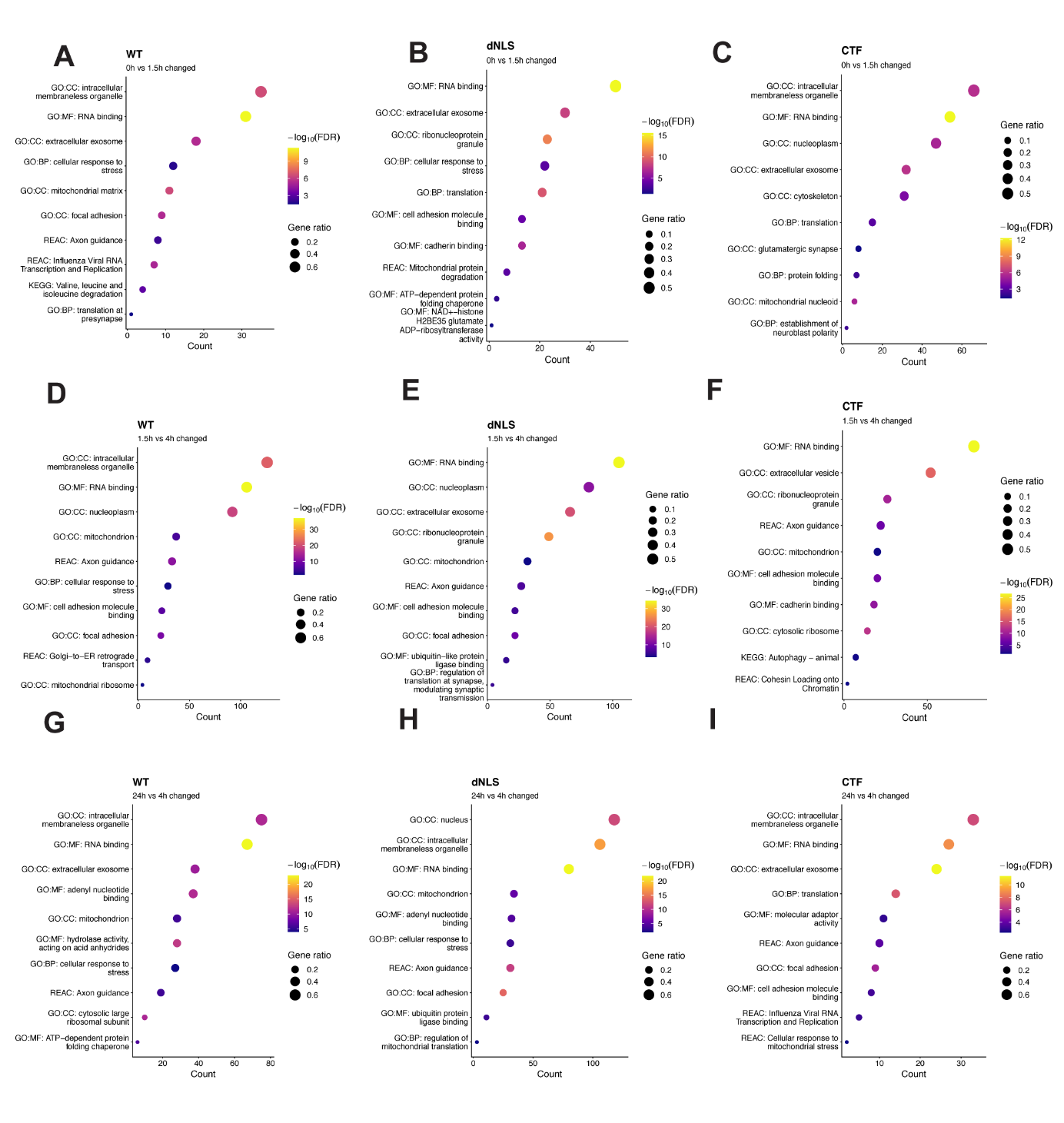

**Figure S4: Gene ontology (GO) enrichment analysis of TDP-43wt, TDP-43dNLS, and TDP-43CTF interactors over the time course.** Significant proteins with p.posthoc < 0.05 and absolute log2FC ≥ 0.5 were used for these analyses**. (A-C)** Terms enriched in unstressed compared to stressed conditions for **(A)** TDP-43wt, **(B)** TDP-43dNLS, and **(C)** TDP-43CTF. **(D-F)** Terms enriched in stressed compared to mid-recovery conditions for **(D)** TDP-43wt, **(E)** TDP-43dNLS, and **(F)** TDP-43CTF. **(G-I)** Terms enriched in mid-recovery compared to recovery for **(G)** TDP-43wt, **(H)** TDP-43dNLS, and **(I)** TDP-43CTF. Complete enrichment results for each comparison is in Supplementary table 2.

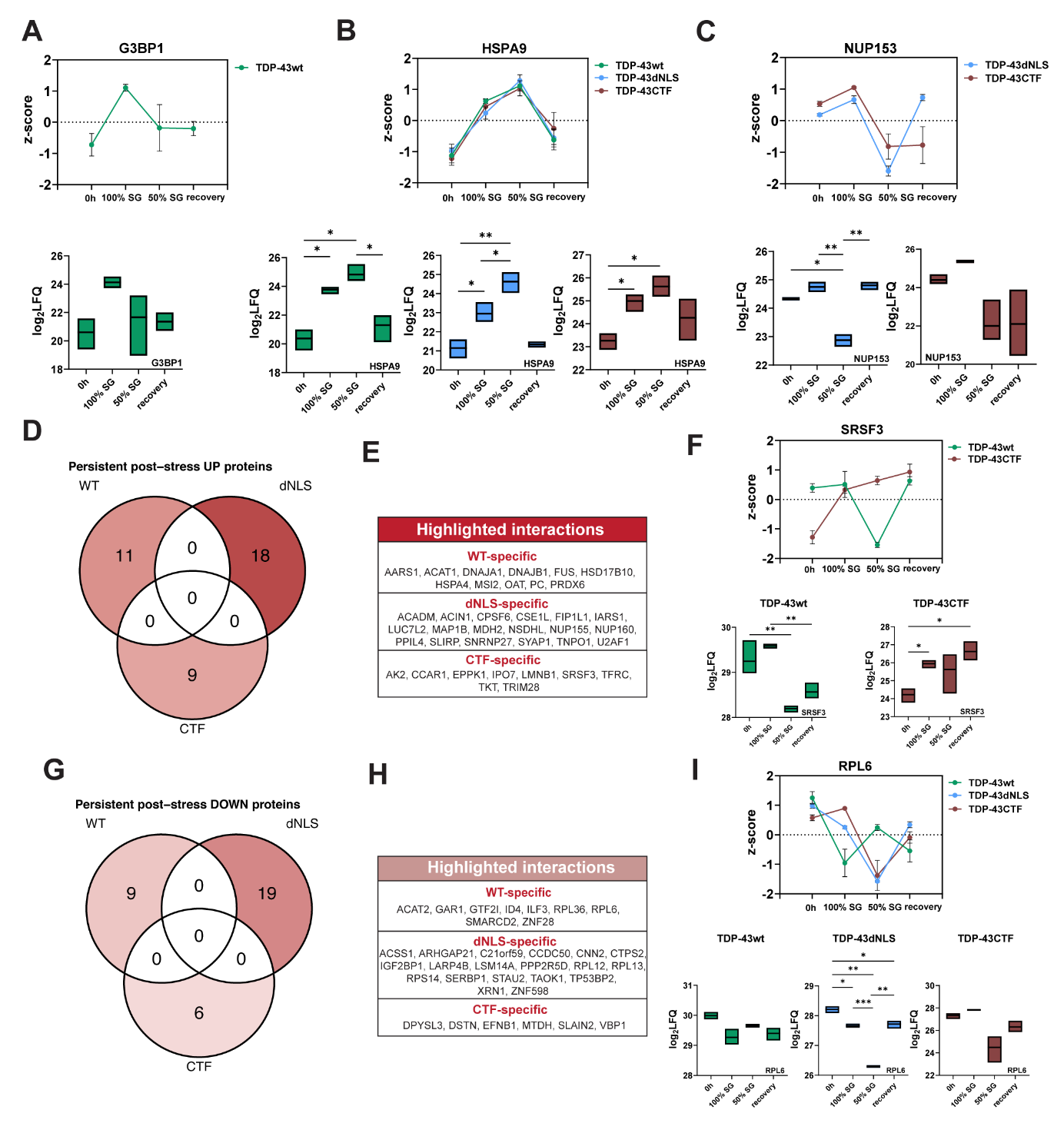

**Figure S5: Persistent post-stress interactors of TDP-43 variants. (A)** Interaction trajectory for interaction between TDP-43wt and stress granule marker G3BP1. Data are presented as z-score mean ± SEM. The corresponding floating bar plot shows log2-transformed LFQ intensity values of G3BP1 over the time course of interaction with TDP-43wt and TDP-43dNLS. Floating bars indicated the min to max range, with a line indicating the mean. Statistical analyses were performed using Welch-ANOVA, followed by a post-hoc test to compare time points in a pairwise manner, *p < 0.05, **p < 0.01, ***p < 0.001. **(B)** Interaction trajectory for interaction between TDP-43wt, TDP-43dNLS, and TDP-43CTF and heat shock protein HSPA9. Data are presented as z-score mean ± SEM. The corresponding floating bar plot shows log2-transformed LFQ intensity values of HSPA9 over the time course of interaction with TDP-43wt, TDP-43dNLS and TDP-43CTF. Floating bars indicated the min to max range, with a line indicating the mean. Statistical analyses were performed using Welch-ANOVA, followed by a post-hoc test to compare time points in a pairwise manner, *p < 0.05, **p < 0.01, ***p < 0.001. **(C)** Interaction trajectory for interaction between TDP-43dNLS, and TDP-43CTF and nucleoporin NUP153. Data are presented as z-score mean ± SEM. **(D)** Venn diagram of upregulated persistent post stress interactors. Proteins were included if they belonged to the persistent post-stress class, were detected in at least 2 of 3 replicates at all four time points, and showed positive change of at least 1.0 at each post-stress time point, and maintained consistent direction between early and late time points. **(E)** Table corresponding to the Venn diagram of upregulated persistent post-stress proteins. **(F)** SRSF3 interaction trajectory with TDP-43 variants over the time course, showing persistent interaction upon stress with TDP-43CTF. Data are presented as z-score mean ± SEM. The corresponding floating bar plot shows log2-transformed LFQ intensity values of SRSF3 over the time course of interaction with TDP-43wt and TDP-43CTF. Floating bars indicated the min to max range, with a line indicating the mean. Statistical analyses were performed using Welch-ANOVA, followed by a post-hoc test to compare time points in a pairwise manner, *p < 0.05, **p < 0.01, ***p < 0.001. **(G)** Venn diagram of upregulated persistent post stress interactors. Proteins were included if they belonged to the persistent post-stress class, were detected in at least 2 of 3 replicates at all four time points, and showed negative change of at least 1.0 at each post-stress time point, and maintained consistent direction between early and late time points. **(H)** Table corresponding to the Venn diagram of upregulated persistent post-stress proteins. **(I)** RPL6 interaction trajectory with TDP-43 variants over the time course, showing persistent loss of interaction upon stress with TDP-43wt. Data are presented as z-score mean ± SEM. The corresponding floating bar plot shows log2-transformed LFQ intensity values of RPL6 over the time course of interaction with TDP-43wt, TDP-43dNLS and TDP-43CTF. Floating bars indicated the min to max range, with a line indicating the mean. Statistical analyses were performed using Welch-ANOVA, followed by a post-hoc test to compare time points in a pairwise manner, *p < 0.05, **p < 0.01, ***p < 0.001.

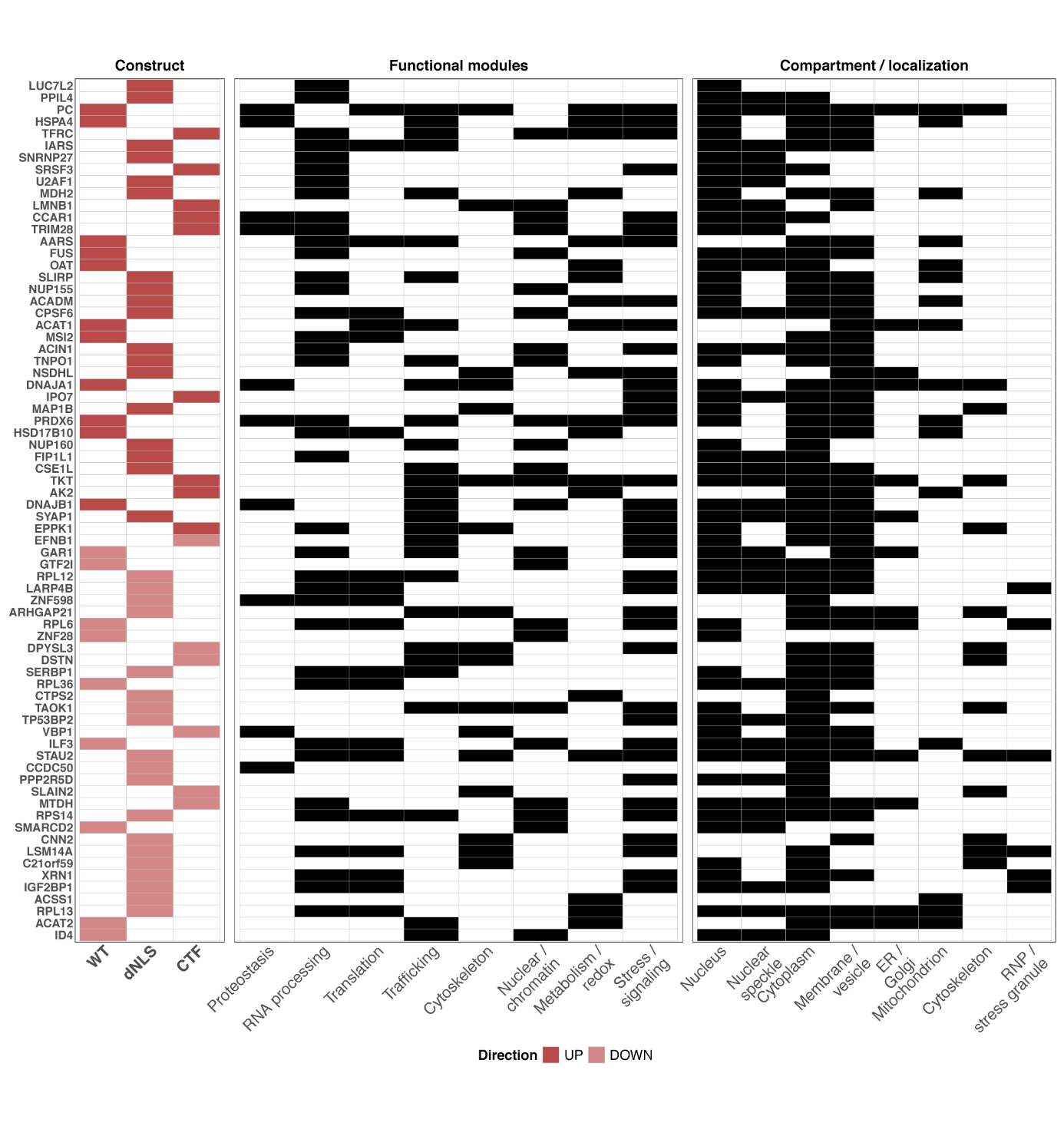

**Figure S6: GO-based descriptive annotation profiling of persistent post-stress TDP-43 interactors.** Upregulated and downregulated candidate proteins associated with persistent interactions gained during stress and persisting post-stress for all constructs were annotated with functional and compartmental features. GO-based annotation panels were generated using descriptive keyword matching of GO terms retrieved through org. HS.eg. db and GO.db.

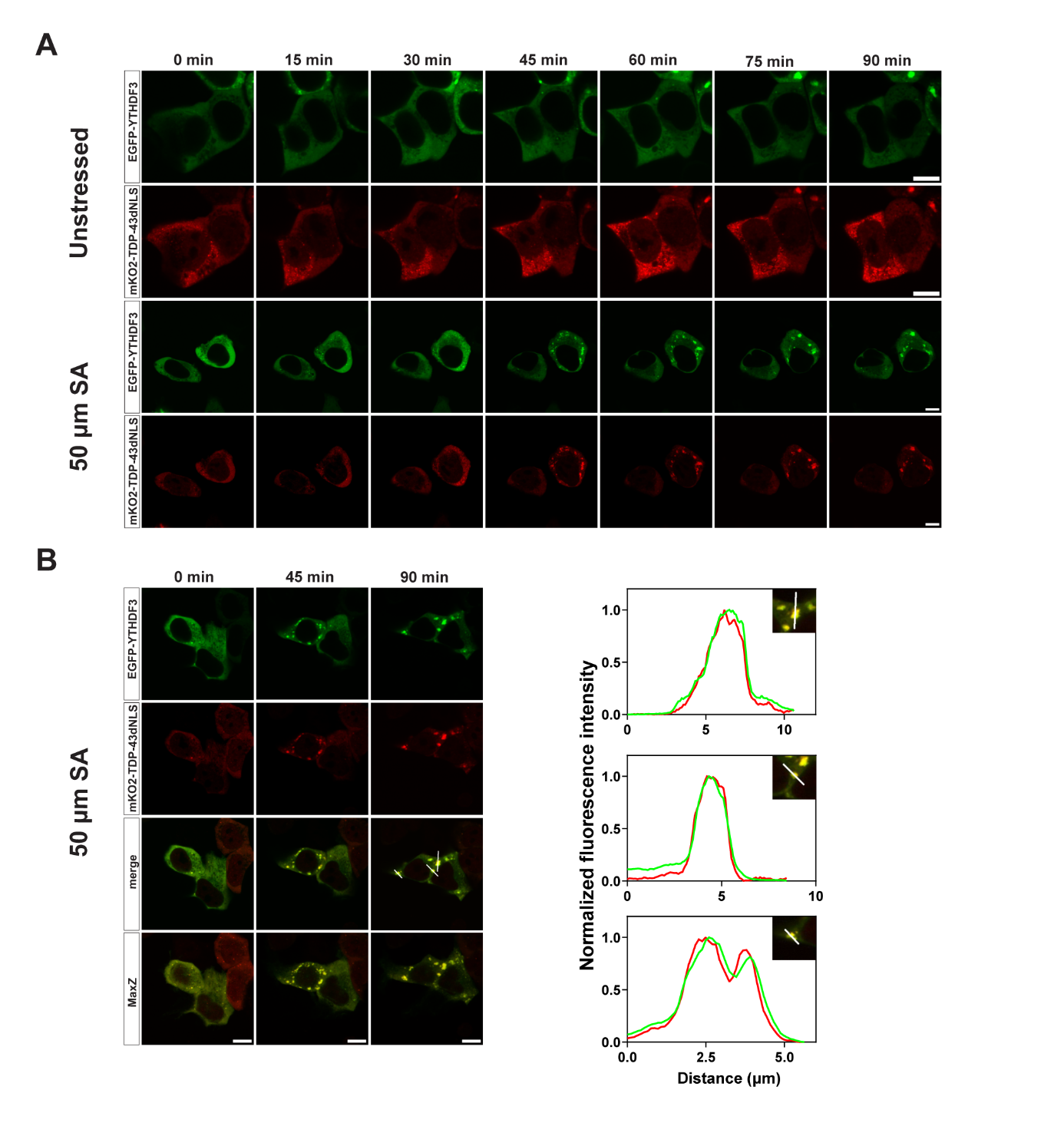

**Figure S7: Live imaging results indicate that TDP-43dNLS granules and YTHDF3 granules appear simultaneously.** Cells were co-transfected with pcDNA5-mKO2-TDP-43dNLS and pT7-EGFP-C1-HsYTHDF3_AB and imaged the following day. **(A)** Unstressed and stressed cells were imaged at 15-min intervals until the final time point at 90 min. TDP-43dNLS granules and YTHDF3 granules appeared simultaneously. **(B)** MaxZ projections of stressed cells at 0-, 45-, and 90-min. Fluorescence intensity profiles for granules at 90 min showed colocalization of these granules. Scale bars, 10 µm.

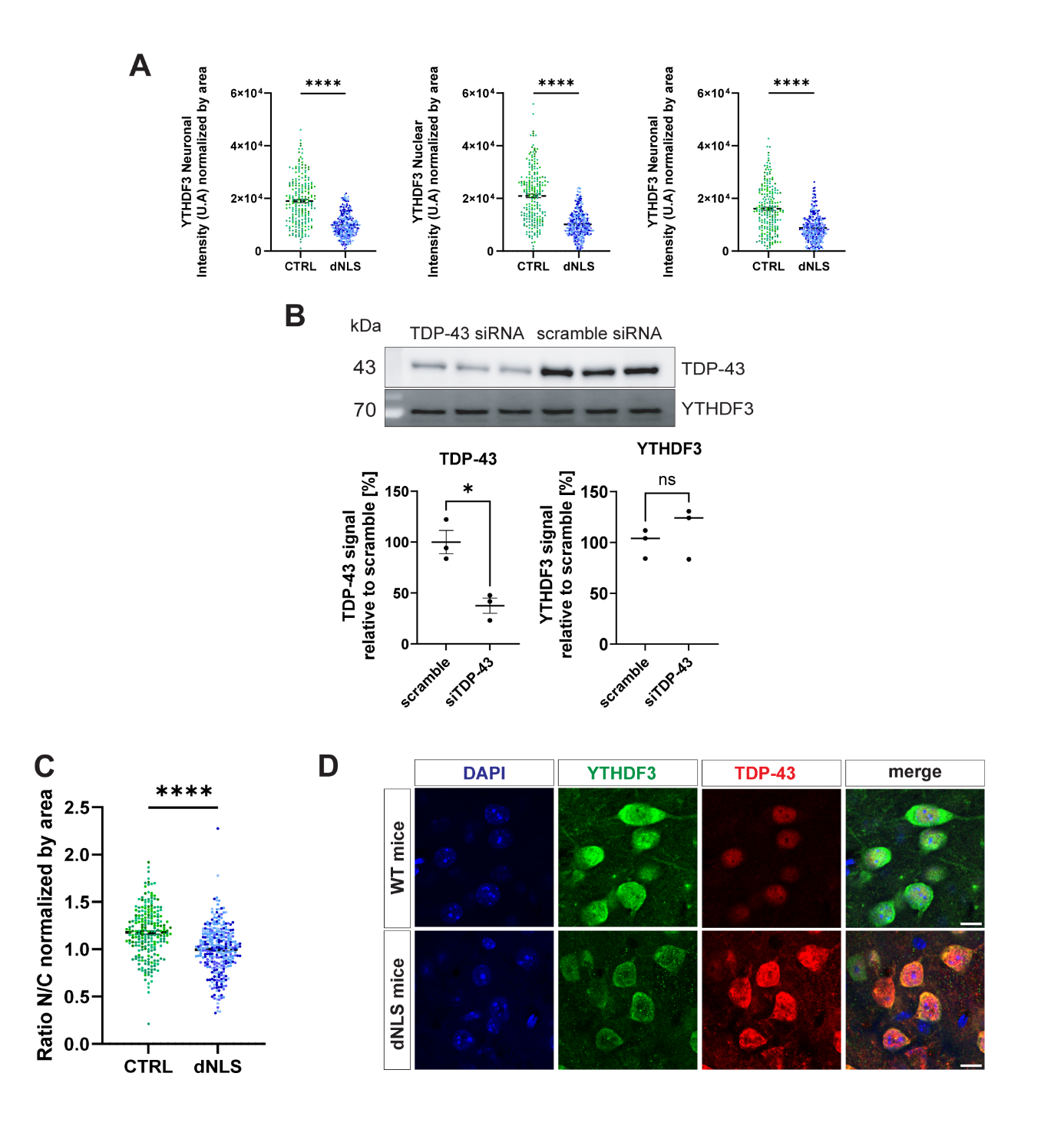

**Figure S8: TDP-43 mislocalization drives YTHDF3 dysregulation. (A)** Graphs showing quantitative analysis of YTHDF3 intensity normalized by the area of whole-neurons, nuclei, and cytoplasm in WT mouse neurons (n = 220) and transgenic TDP-43dNLS mouse neurons (n = 298). **(B)** Western blot analysis of KD TDP-43 HEK293 cells. Western blots were probed for TDP-43 and YTHDF3. The experiment was performed in three biological replicates. Graphs show quantitative analysis of TDP-43 signal relative to scramble and YTHDF3 signal relative to scramble. Statistical analyses were performed using a paired two-tailed t-test; ns, not significant, *p < 0.05, data are shown as mean ± SEM. **(C)** Quantification of the nuclear-cytoplasmic ratio normalized by area in WT mouse neurons (n = 220) and transgenic TDP-43dNLS mouse neurons (n = 298). Brain tissue from four mice per group was analyzed; different shades of blue and green correspond to different animals. Statistical analyses were performed using an unpaired two-tailed t-test; ****p < 0.0001; data are shown as mean ± SEM. **(D)** Confocal images of large mouse neurons from mice expressing TDP-43wt and transgenic mice expressing TDP-43dNLS, immunolabelled with anti-TDP-43 antibody (red) and anti-YTHDF3 antibody (green). DNA was labelled with 4,6-diamidino-2-phenylindole (DAPI) (blue). Scale bars, 10 µm.

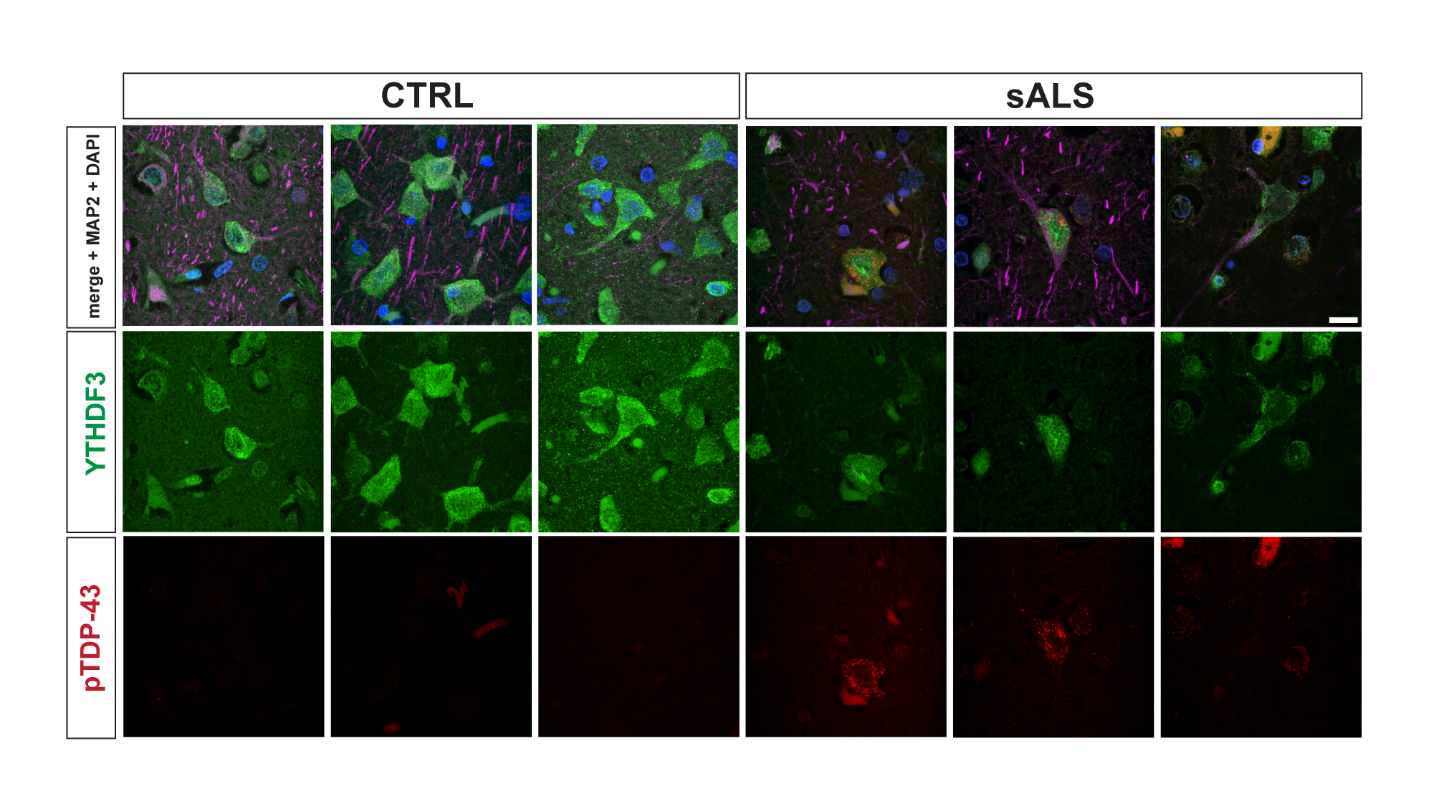

**Figure S9: Interaction between TDP-43 and YTHDF3 in ALS/FTD pathology.**  Additional examples of human motor cortex from representative controls and sporadic ALS patients immunolabelled with anti-TDP-43 antibody (red), anti-YTHDF3 antibody (green) and anti-MAP2 (violet). DNA was labelled with 4,6-diamidino-2-phenylindole (DAPI) (blue). Images are presented as z-stack projections. Scale bar, 10 µm.

**Supplemental Tables**

**Table S1: Proteomics analysis and Statistical Data, related to Figures 2, 3, 6 and Figures S3 and S5.** This table provides the complete computational output from the proteomics analysis, including z-scores, On/Off analysis, fold changes and the corresponding statistical analyses.

**Table S2: Gene Ontology Analysis Across All Time Points and TDP-43 Constructs, related to Figure S4.** Gene Ontology (GO) analysis was performed for proteins enriched at different time points during the stress and recovery phases for TDP-43WT, TDP-43dNLS, and TDP-43CTF. In the main manuscript, we present statistically significant GO terms associated with neurodegeneration, whereas this table provides the complete set of statistically significant GO terms identified in the analysis.

**Table S3: Gene Ontology Analysis of Late-Recovery and Persistent TDP-43 Interactors, related to Figures 4-5 and Figures S5-S6.** GO-based descriptive annotation profiling was performed for late-recovery-associated and persistent post-stress interactors of TDP-43wt, TDP-43dNLS, and TDP-43CTF. This table presents all GO terms associated with these proteins. In the main manuscript, related GO terms are grouped into broader common categories to improve clarity and facilitate interpretation.

**Table S4: Mouse and Human Pathology-Related Information, related to Figure 7 and Figures S8-S9.** This table provides information on the mice used in this study, as well as clinical and pathology-related information for all human ALS and control (CTRL) cases included in the study.

**Table S5: Primers, Antibodies, and Buffer Compositions.** This table provides information on all primers used for molecular cloning, the precise compositions of buffers used for pull-down experiments, and the antibodies and corresponding dilutions used for different experimental applications described in the manuscript.

**Table S6: Raw data from knockdown, live-imaging, line-plots and mouse brain analyses, related to Figure 6-7 and Figures S7-S8**. This file contains the raw data underlying the graphs and line plots presented in the manuscript, including data from knockdown experiments, live-imaging experiments, and mouse brain analyses.
